# Integrative vasculogenesis unifies distinct endothelial sources in the developing lung

**DOI:** 10.64898/2026.05.08.723805

**Authors:** Miram Meziane, Mackenzie P.H. Litz, Prashant Chandrasekaran, David Frank, Pulin Li

**Affiliations:** Department of Biology, Massachusetts Institute of Technology, Cambridge MA, 02139; Computational and Systems Biology Program, Massachusetts Institute of Technology, Cambridge MA, 02139; Whitehead Institute for Biomedical Research, Cambridge MA, 02139; Department of Pediatrics, Division of Cardiology, University of Pennsylvania, Children’s Hospital of Philadelphia (CHOP), Penn-CHOP Lung Biology Institute, Penn Cardiovascular Institute, CHOP Cardiovascular Institute, Philadelphia, PA 19104

## Abstract

How endothelial cells from distinct developmental sources are integrated into a single continuous vascular system remains unresolved. Here, using the developing mouse lung, we identify a mesenchymal progenitor population that generates endothelial cells *de novo* and incorporates them into the expanding vasculature through a mechanism we term integrative vasculogenesis. Genetic lineage tracing shows that these progenitors contribute directly to the pulmonary endothelium, defining a source distinct from endothelial cells of the major vessels. Live imaging and single-cell tracking reveal that newly specified angioblasts exhibit high motility, dispersing through stochastic migration before integrating into pre-existing vascular networks. Cell ablation demonstrates that pre-existing networks are required to support the migration, proliferation and survival of nascent endothelial cells. Integrative vasculogenesis is thus distinct from classical vasculogenesis and angiogenesis, providing a framework for how endothelial populations of different origins are assembled into a functional circulatory system.

## Introduction

The rapid growth and morphological changes during organogenesis require the coordinated assembly of a functional vascular system to ensure continuous delivery of oxygen and nutrients. This, in turn, depends on the integration of organ-specific vasculature with the systemic circulation. However, the origin of organotypic endothelial cells and how they assemble into a single continuous network remain highly debated^1^.

Central to this debate is whether organotypic endothelial cells originate from within the organ or are supplied from external sources. In the liver and lung, major vessels are connected to the heart, and progenitors within the heart field contribute to their formation^2–4^. In parallel, organ-restricted endodermal or mesodermal progenitors have been proposed to generate endothelial cells locally^5–10^. Yet, key endothelial progenitors, such as ETV2+ angioblasts, were never identified within developing organs, even though they have been observed outside visceral organs^11^. These observations raised the possibility that multiple spatially and temporally distinct endothelial sources could co-exist during organogenesis.

If multiple endothelial populations contribute to organ vascularization, how they are assembled into a single, continuous network cannot be explained by current models of blood vessel formation. Classic vasculogenesis describes the *de novo* formation of vascular networks from angioblasts, whereas angiogenesis expands existing vessels into surrounding tissues^12–15^. However, these models do not account for how independently arising endothelial populations are subsequently integrated into a unified circulatory system.

The lung provides a tractable system to address this problem. Although multiple endothelial sources for the lung have been proposed based on lineage tracing, the broad expression of Cre drivers and temporal labeling windows obscure their independence^3,4,16^. Furthermore, analysis based on fixed tissues have led to conflicting interpretation of vascular assembly^17–19^. Resolving these issues requires direct identification of endothelial progenitor sources and tracking their behavior in live tissues.

Here, we identified a distinct *Wnt2+Flk1+* mesenchymal progenitor population that generates endothelial cells *de novo* that stably integrate into the pulmonary vasculature. Rather than forming a separate vascular plexus, newly specified angioblasts migrate and incorporate directly into pre-existing networks, a process we define as integrative vasculogenesis. The migration, proliferation and survival of nascent endothelial cells relies on existing networks, which undergo angiogenesis concurrently, providing a framework for how distinct endothelial populations are unified into a continuous circulatory system, with implications for engineering organotypic vasculature *in vitro*.

### Respiratory mesenchymal progenitors are spatiotemporally diverse

To capture the initial vascularization events of the lung in an unbiased fashion, we performed scRNA sequencing on lungs and adjacent tissues from 36 E10-E12.5 embryos (**Fig. 1A**). After quality control and filtering, a total of 12,470 high-quality cells were recovered, the majority of which corresponded to mesenchymal cells, defined by their expression of *Pdgfra* and *Foxf1*, in addition to endothelial cells, endodermal cells, neural crest and neurons (**Figs. 1B, Extended Data Fig. 1A**)^20^. After filtering out mesenchymal cells that represent other organ identities, we revealed heterogeneous *Tbx4+Tbx5+* respiratory mesenchymal cells at this early stage of lung development (**Figs. 1C, D, Extended Data Fig. 1B**). In addition to three known committed mesenchymal populations, the mesothelium, *Acta2+* smooth muscle cells, and *Sox9+Col2a1+* chondrocyte progenitors, we discovered a *Smoc2+* population localized immediately adjacent to lung epithelia (**Figs. 1D, Extended Data Figs. 1C, 2A,B**)^21^.The spatial domain occupied by this population at E10.5 resembled that of airway smooth muscle cells at E11.5, suggesting the possibility of *Smoc2+* cells being smooth muscle progenitors (**Extended Data Figs. 2A, B**). Notably, both smooth muscle and chondrocyte progenitors were primarily detected in the anterior buds and proximal airways, suggesting a delayed cell fate commitment among mesenchymal progenitors in the posterior buds (**Extended Data Figs. 2C, D**).

**Figure 1.**
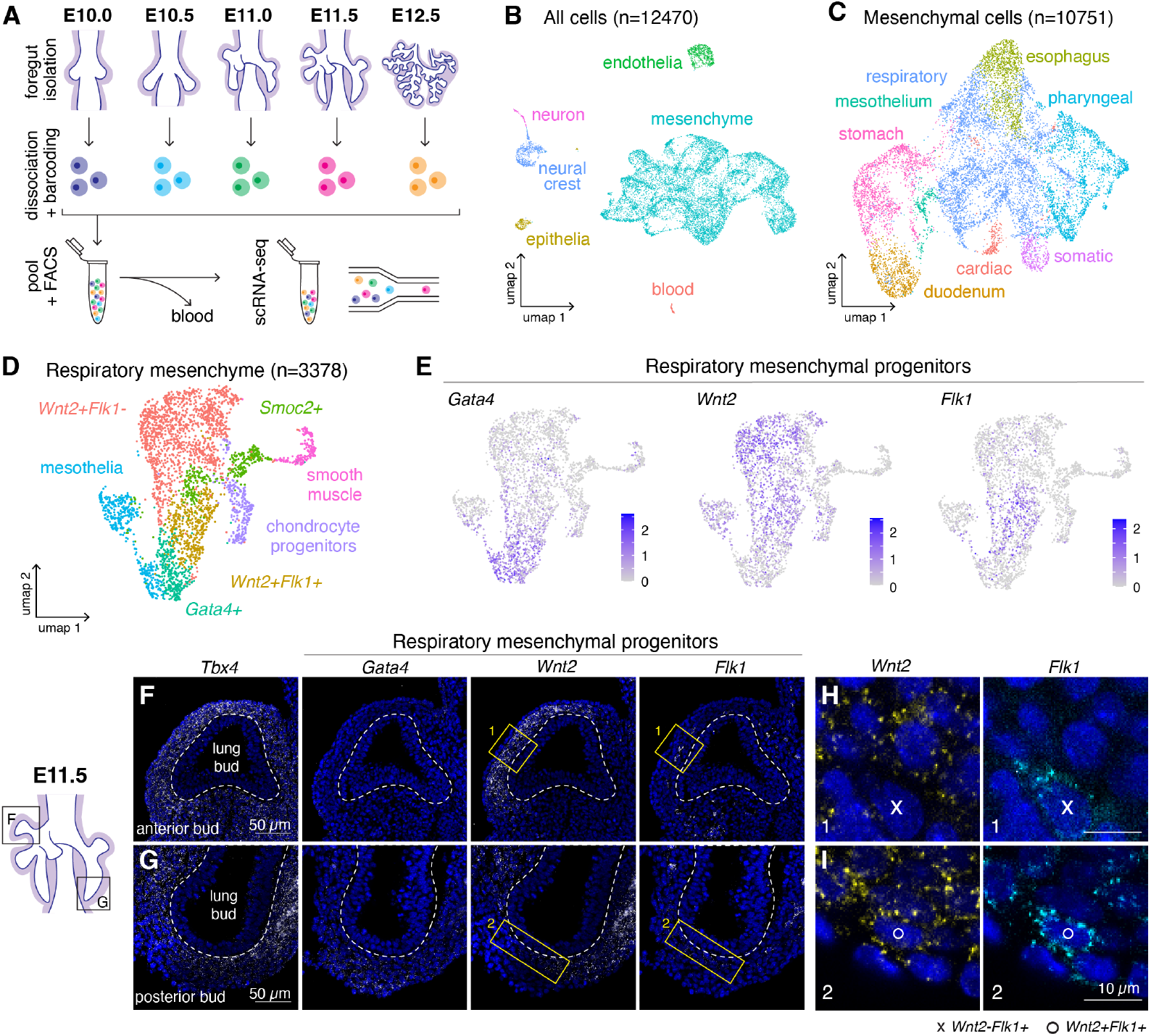
Mesenchymal progenitor heterogeneity in the embryonic lung. **A**. Schematic of scRNA-seq experiment. Embryos were isolated, staged, dissociated, and barcoded accordingly. Samples were pooled for sequencing after removal of blood cells by FACS. **B**. UMAP representation of all cells collected after quality control, annotated by cell type. **C**. UMAP representation of only mesenchymal cells (*Pdgfra+Foxf1+* and *Pdgfra+Foxf1-*), annotated by organ. **D**. Unsupervised clustering of respiratory mesenchyme (*Foxf1+Tbx4+Tbx5+)* and annotated by cell type or by differentially expressed genes. E. Expression of progenitor markers across the respiratory mesenchyme. **F-G**. Spatial distribution of respiratory mesenchymal populations by hybridization chain reaction (HCR)^47^ in an anterior (**F**) and posterior (**G**) bud of an E11.5 lung. **H-I**. Inset of regions 1 (**H**) and 2 (**I**).

Furthermore, we recovered three populations that closely resembled uncommitted mesenchymal progenitors, each with distinct spatial organization (**Figs. 1D-G, Extended Data Figs. 2E, F**). A transient *Tbx4+Gata4+* population, which was previously implicated in lobe formation, was most prominent on the ventral side of the E10.5 lung and later became maintained only in the posterior-most buds adjacent to the mesothelium at E11.5 (**Figs. 1F,G, Extended Data Figs. 2E, F**)^22^.While remaining cells all expressed the lung mesenchymal progenitor marker *Wnt2* (**Fig. 1E**)^3,16,23^, *Wnt2+* cells could unexpectedly be further subdivided by the presence or absence of *Flk1* (**Fig. 1D,E**). At E10.5, *Wnt2+Flk1-* cells were distributed dorsolaterally in a pattern opposing that of *Gata4* and became concentrated at distal tips in both the anterior and posterior buds by E11.5 (**Figs. 1F, Extended Data Figs. 2E, F**). In contrast, *Wnt2+Flk1+* cells were observed exclusively in the posterior-most lung buds across all timepoints examined (**Figs 1F-I, Extended Data Figs. 2E, F**). Altogether, our data demonstrates a previously unappreciated temporal and spatial heterogeneity within the respiratory mesenchyme and highlighted that the posterior buds display delayed mesenchymal differentiation states relative to anterior buds.

### An angioblast population exists within the *Wnt2+Flk1+* domain of the lung

The *Wnt2+Flk1+* population was surprising: these cells simultaneously maintained lung mesenchymal progenitor identity (*Tbx4, Tbx5, Wnt2, Fgf10*) and endothelial potential (*Flk1*), yet were devoid of mature endothelial markers (*Cdh5, Pecam1*) (**Extended Data Figs. 3 A, B**). Although *Flk1+* progenitors in the lateral plate mesoderm and placenta are known to produce endothelial cells^24^, such mesodermal progenitors with endothelial potential were never directly observed in developing organs after having obtained organ identity. Therefore, we hypothesized that this *Wnt2+Flk1+* population might reflect an organ-specific mesenchymal progenitor that generates angioblasts to vascularize the lung.

To confirm that this population is distinct from differentiated endothelial cells and harbor angiogenic potential, we stained the lung with markers representing different stages of endothelial commitment and differentiation (**Fig. 2A-H**).

**Figure 2.**
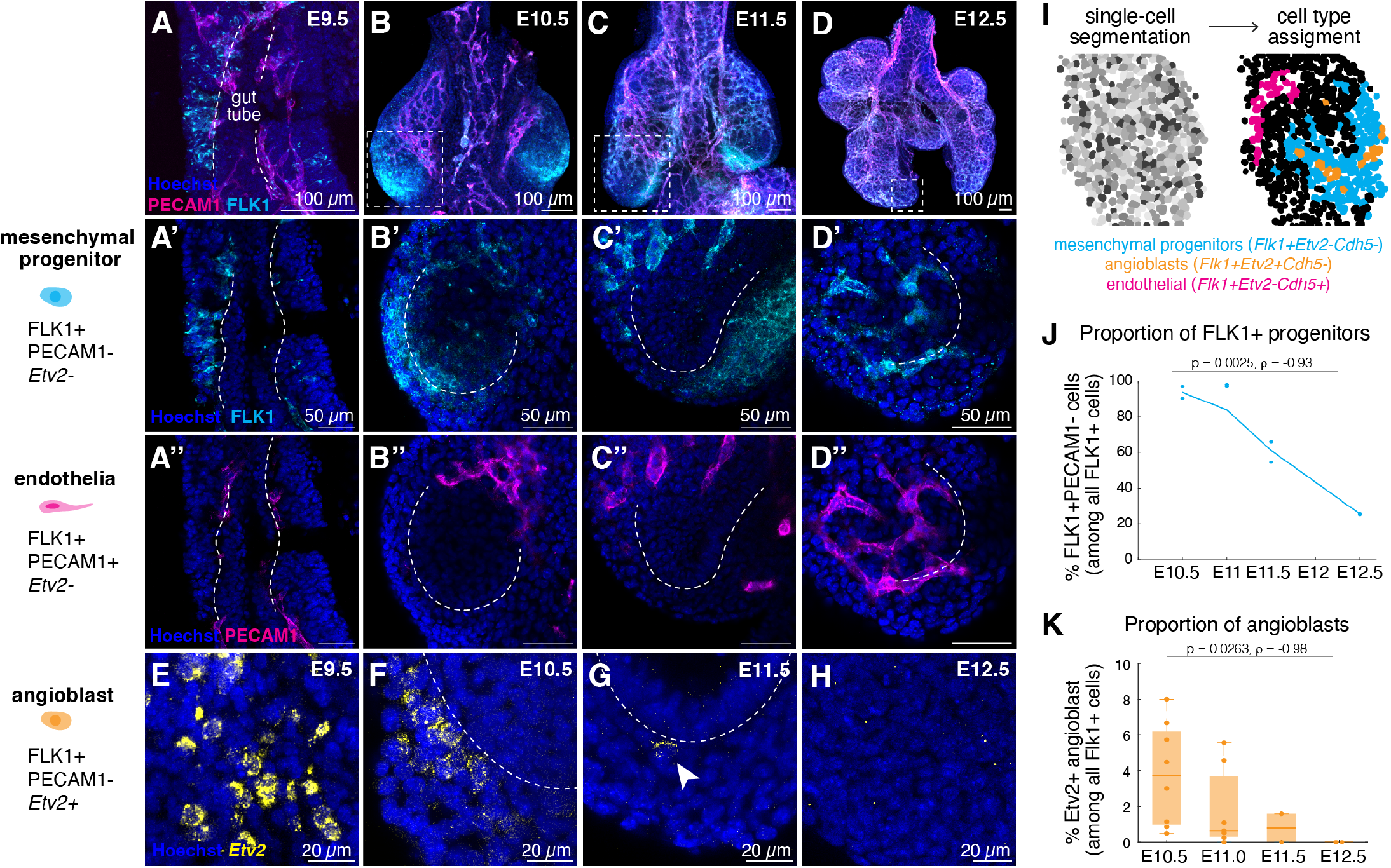
*Wnt2+Flk1+* defines a posterior domain with *de novo* vascularization potential. **A-D**. Whole mount antibody staining and imaging of E9.5 foregut and E10.5-E12.5 lung stained for FLK1 and endothelial marker PECAM1. **A’-D’’**. Optical section of the foregut (**A’**,**A’’**) or posterior-most lung buds (**B’-D’’**), corresponding to the boxed regions in **B-D**. Images are split by channel to highlight the spatial differences in expression of FLK1 (**A’-D’**) and PECAM1 (**A’’-D’’**). **E-H**. Presence of *Etv2+* angioblasts in the foregut (**E**) and posterior-most lung buds (**F-H**). Images showing HCR signals in a single optical section. **I**. Example masks generated by cell segmentation of nuclei (Hoechst) in whole-mount posterior buds before (*left*) and after (*right*) cell-state assignment. **J**. Quantification of the proportion of FLK1+PECAM1-cells among all FLK1+ cells in posterior bud FOV. 2 buds for each time point were used, with 3000-6000 cells profiled at each time point. A two-sample t-test was performed between E10.5 and E12.5 quantification to assess significance. Significance of the trend (ρ) was assessed by Spearman’s correlation. **K**. Quantification of proportion of *Etv2*+ cells among all *Flk1+* cells in posterior bud FOV. Between 2 and 8 buds were examined for each time point, with a minimum of 2000 cells segmented per bud. Two-sample Kolmogorov-Smirnov test was performed between E10.5 and E12.5 quantification to assess significance. Significance of the trend (ρ) was assessed by Spearman’s correlation.

FLK1+PECAM1-progenitors and FLK1+PECAM1+ differentiated endothelial cells were initially intermingled adjacent to the gut endoderm at E9.5, and became spatially separated by E10.5, with progenitors enriched only at the avascular tips of the lung buds (**Fig. 2A, B**). This progenitor domain persisted in the posterior-most buds of the E11.5 lung, while anterior buds became surrounded by a PECAM1+ vascular networks (**Fig. 2C**). By E12.5, the progenitor domain disappeared, coinciding with a complete vasculature surrounding the posterior lung buds (**Fig. 2D**). *Etv2*+ angioblasts, which we detected in a small fraction of the endothelial cluster in our scRNA-seq data (**Extended Data Fig. 3B**), followed the same spatiotemporal pattern as FLK1+PECAM1-progenitors, peaking at E10.5 within the FLK1+ domain and disappearing by E12.5 (**Figs. 2E-H, Extended Data Fig. 3C**). Quantification of relative abundance in the posterior lung buds by single-cell segmentation confirmed a consistent decrease in the proportion of both FLK1+PECAM1-progenitors and *Etv2*+ angioblasts among all FLK1+ cells between E10.5-E12.5 (**Fig. 2I-K**). Together, these suggest that *Wnt2+Flk1+* mesenchymal progenitors provide a transient source of angioblasts that produce endothelial cell *de novo* in the developing lung.

### *Wnt2+ Flk1+* progenitors stably contribute to diverse pulmonary endothelial cells

To experimentally validate the potential of *de novo* vascularization in the developing lung, we performed an endothelial ablation-recovery experiment on lung explants. Differentiated endothelial cells uniquely rely on vascular endothelial growth factor (VEGF) signaling for their survival^25–27^. We treated cultured E10.5 lungs for 24 hrs with SU5416, a chemical antagonist of the receptor VEGFR2 (FLK1) and assessed endothelial survival by probing for expression of *Cdh5*, which labels differentiated endothelial cells similarly to PECAM1 (**Extended Data Fig. 3B**). The treatment completely ablated *Flk1+Cdh5+* endothelial cells but spared the *Wnt2+Flk1+Cdh5-* progenitor population, with no impact on the overall structure of the lung buds (**Extended Data Fig. 4A**)^28,29^. We then allowed the chemically-ablated lungs to recover *ex vivo* for an additional 24 hrs after removal of the drug. *Flk1+Pecam1+Cdh5+* endothelial cells appeared only in the posterior buds, confirming it being a privileged region for *de novo* endothelial generation (**Fig. 3B, Extended Data Fig. 4B**). In contrast, explants kept in the drug during this time were unable to generate new endothelial cells (**Fig. 3C**).

**Figure 3.**
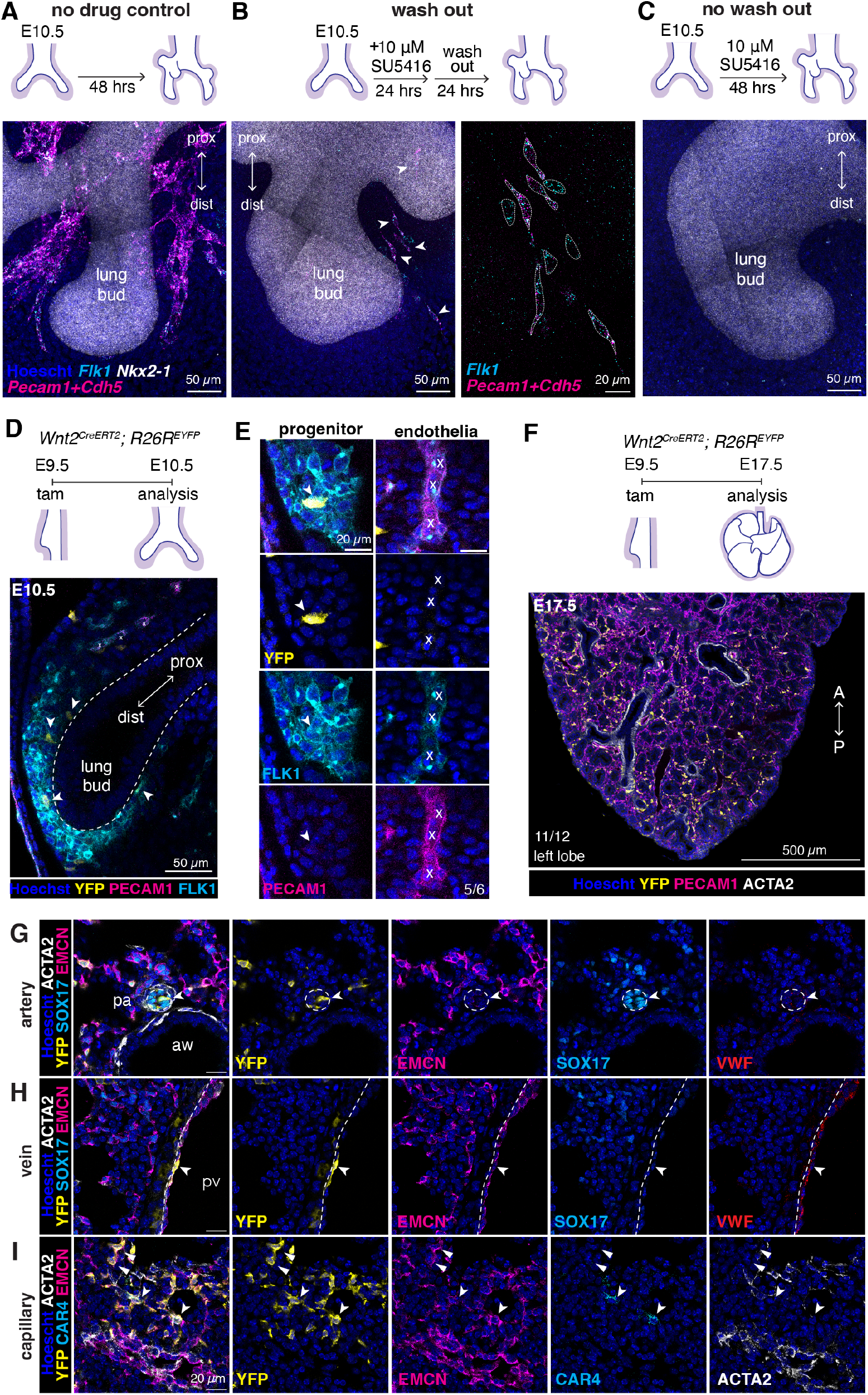
*Wnt2+ Flk1+* progenitors stably contribute to diverse pulmonary endothelial cells. **A-C**. Ablation recovery experiments on E10.5 lung explants. Distribution of endothelial cells after 48 hrs of culture with no drug treatment (**A**), after 24 hrs of chemical ablation followed by 24 hrs of recovery without the drug (**B**), or in the continued presence of SU5416 (**C**). Inset in (**B**) shows scattered *Flk1+* cells that are either progenitors (*Pecam1-Cdh5-)* or differentiated endothelial cells (*Pecam1+Cdh5+)* in the posterior-most bud. **D-I**. Lineage tracing confirmed stable contribution of mesenchymal progenitors to diverse endothelial cell types. Representative image of E10.5 lung isolated 24 hrs post tamoxifen injection, stained for YFP, FLK1 and PECAM1, to validate the labeling strategy (**D**). FLK1+PECAM1-progenitors were labelled, whereas FLK1+PECAM1+ endothelia were unlabeled, among 5 out of 6 E10.5 lungs from two litters examined (**E**). **F**. Schematic of tracing *Wnt2*+ cell fates long-term and representative section of the left lobe of an E17.5 lung post tamoxifen administration. 11 of 12 samples from 2 litters examined had YFP+ endothelia. **G-I**. Sections of E17.5 lung demonstrating labelled subtypes of endothelial cells, including artery (**G**), and vein (**H**) and capillary (**I**) cells. aw: airway. pa: pulmonary artery. pv: pulmonary vein.

To directly test if *Wnt2+* respiratory mesenchymal progenitors stably contribute to the pulmonary vasculature, we performed lineage tracing using the *Wnt2*^*CreERT2*^; *R26R*^*EYFP*^ mice^3,30^. We administered tamoxifen as a pulse on day E9.5, when *Wnt2+* mesenchymal progenitors have gained respiratory identity^31^. Despite the low labeling efficiency that was consistent with previous studies, both FLK1+ and FLK1-mesenchymal cells were labelled at E10.5, with FLK1+ labeled cells concentrated in the posterior of the lung buds (**Figs. 3D, E**)^3,4^. Importantly, we did not observe any labelled FLK1+PECAM1+ endothelial cells in 5 of the 6 samples examined (**Fig. 3E**), with 1 of 6 samples displaying YFP+ endothelia.

To identify the cell types derived from *Wnt2+* cells labelled at E9.5, we stained the lung at E17.5 and found that the progenies of labeled *Wnt2+* cells were broadly distributed in the lung stroma, consistent with previous studies (**Fig. 3F**)^3,4^. Importantly, of 12 YFP+ samples examined, we recovered 11 with labelled endothelial cells scattered across the E17.5 lung, representing the artery (SOX17+EMCN-), vein (SOX17-EMCN+VWF+), and capillary (EMCN+) (**Fig. 3G-I**). We did not observe labeled lymphatic endothelial cells (NRP2+,PECAM1-), suggesting their separate origin (**Extended Data Fig. 5A**)^32,33^. While most labelled capillary cells were CAR4-, we observed rare instances of labelled CAR4+EMCN+ capillary cells, suggesting that mesenchymal progenitors can give rise to this specialized capillary cell which first appears at E17.5 (**Fig. 3I**)^34,35^. Similar results were obtained when tamoxifen was injected at E10.5, whereas injection at E12.5 failed to generate labeled endothelial cells, even though other stromal cell types were still labeled (**Extended Data Figs. 5B-E**). Together, these results demonstrate the transient *Wnt2+Flk1+* progenitor pool supplies endothelial cells *de novo* after initial lung bud formation and stably contributes to the vasculature.

Although the posterior *Wnt2+Flk1+* mesenchymal progenitors supply endothelial cells within the lung, *Wnt2+* mesenchymal cells surrounding the heart field and anterior foregut prior to lung specification at E8.5 also contribute to the pulmonary endothelium, referred to as cardio-pulmonary progenitors (CPPs)^3,4^. To compare our *Wnt2+Flk1+* lung mesenchymal cells with those of the CPP, we examined the spatiotemporal distribution of *Flk1* and CPP markers *Wnt2, Isl1* and *Gli1*. We found that CPPs were devoid of *Flk1* expression at E8.5 (**Extended Data Fig. 6A**). Reciprocally, we found that while *Wnt2+Flk1+* cells expressed *Gli1*, which is also broadly expressed in the lung mesenchyme, these cells displayed little to no expression of *Isl1* (**Extended Data Figs. 6B, C**). We could detect cells on the anterior and ventral region of the lung displaying expression of all 3 CPP markers, however these did not express *Flk1* and importantly, were found opposite the *Wnt2+Flk1+* progenitors, localized distally and on the dorsal side of the lung (**Extended Data Fig. 6C**). Therefore, the *Wnt2+Flk1+* population is spatially distinct from CPPs, although we cannot completely rule out the possibility that the *Wnt2+Flk1+* population is derived from the CPP during lung specification.

### Angioblasts extend existing networks through integrative vasculogenesis

The spatial segregation between pre-existing vascular networks and the *Wnt2+Flk1+* progenitor pool raises the question of how *de novo* generated endothelial cells stably and seamlessly integrate into the pre-existing vasculature of the lung. The classic model of *de novo* vascularization is vasculogenesis, in which angioblasts assemble into a primitive network that later gets remodeled as the cells differentiate into endothelial cells, as reported in the yolk sac and dorsal aorta^36,37^.

However, we did not observe the assembly of disparate networks. Instead, through live imaging of E10.5 lung isolated from Tg(*Flk1:GFP*) mice *ex vivo*, we observed continuous incorporation of differentiating angioblasts into existing networks over the course of 30 hrs (**Fig. 4A, Supplementary Video 1**)^38^. By tracking individual cells over time, we identified three populations defined by their GFP expression dynamics: cells expressing a constantly low or high level of GFP (FLK1_low and FLK1_high), as well as cells that switched from low to high expression (FLK1_switch) (**Fig. 4B**). Staining the lung with cell fate markers confirmed these populations correspond to *Flk1+* mesenchymal progenitors (FLK1_low), differentiated endothelial cells (FLK1_high) and differentiating angioblasts (FLK1_switch, corresponding to intermediate GFP levels) (**Fig. 4C, Extended Data Fig. 7A**). The cell morphology further confirmed the states of GFP+ cells, with FLK1_low cells exhibiting a rounded, progenitor-like morphology and FLK1_switch cells gradually adopting an elongated shape reminiscent of differentiated endothelial cells (**Figs. 4D, E, Supplementary Video 2**).

**Figure 4.**
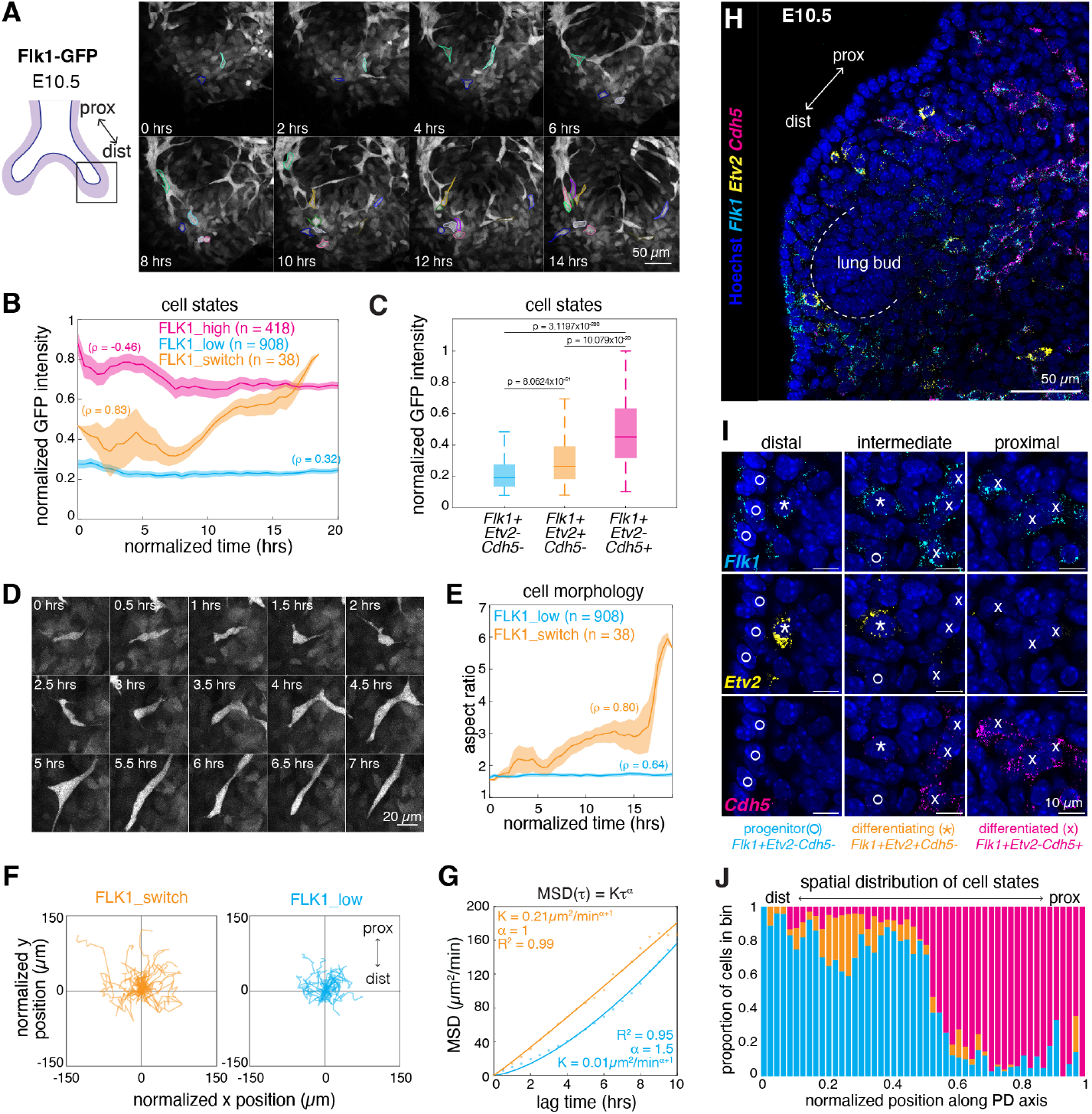
Angioblasts extend existing networks through integrative vasculogenesis. **A**. Montage of timelapse imaging on E10.5 FLK1-GFP explants. Outlined cells represent differentiating (FLK1_switch) cells. Each color represents a single cell over time. **B**. GFP expression dynamics representing different cell states. Trajectories were aligned based on either the start time of the trace (for FLK1_high and FLK1_low trajectories) or the point at which FLK1_switch traces transition from low to high GFP expression. Significance of the trend (ρ) was assessed by Spearman’s correlation. **C**. GFP expression levels among different *Flk1+* cell populations in E10.5 FLK1-GFP mouse lungs. Populations were identified based on multiplexed HCR (Fig. S7A). n = 1925 cells across two independent samples. A two-sample t-test was performed between populations to assess significance. **D**. Montage of a representative FLK1_switch cell, showing dynamic changes in cell morphology. **E**. Quantification of cell morphology dynamics across all FLK1_switch cells vs FLK_low cells over time. Significance of the trend (ρ) was assessed by Spearman’s correlation. **F**. Overlay of all FLK1_switch and 50 randomly selected FLK1_low traces, normalized to the cell’s initial position at t_norm_ = 0 hrs. **G**. Mean squared displacement (MSD) of FLK1_switch and FLK1_low trajectories during the first 10 hrs of each trajectory. **H**. Optical section of E10.5 lung stained for endothelial markers. **I**. Representative images of different FLK1+ populations along the proximal-distal axis. **J**. Distribution of FLK+ populations along the normalized proximal-distal axis, split into 50 evenly distributed bins. n = 3707 cells across 3 independent samples.

Upon commitment to an endothelial fate, differentiating cells became more migratory, searching for stable contacts with other endothelial cells. The FLK1_switch cells emerged asynchronously over time, and exhibited a more rapid search of their local environment than FLK1_low cells (**Figs. 4F, Extended Data Figs. 7B, C**). The migration behavior appeared to be largely random, rather than directionally guided, suggesting that increased motility is an intrinsic feature of differentiating angioblasts (**Fig. 4G, Extended Data Figs. 7C**). Importantly, the search allowed FLK1_switch cells to approach differentiated endothelial cells that are located more proximally and make contact, before eventually incorporating into existing networks (**Extended Data Figs. 7D, E**).

Adhesion-mediated cell-cell interactions likely “captured” differentiating cells to prevent them from moving away, which is consistent with the expected accumulation of endothelial adhesion molecules during angioblasts differentiation. Indeed, reconstructing the differentiation trajectory from *Wnt2+Flk1+* progenitors to differentiated endothelial cells in our scRNA-seq data revealed a gradual upregulation of adhesion molecules after the pulse-like expression of *Etv2* and commitment to an endothelial fate (**Extended Data Fig. 8**)^39^. Altogether, these data suggests that the directional expansion of vascular network is a result of a random search of differentiating endothelial cells originating from the distal end, which are likely captured by the existing network in the opposing, proximal end, although our results do not completely exclude the possibility of small bias in migration direction.

The spatial distribution of the three populations further confirmed this migration-capture model. By using multiplexed HCR, we assigned progenitor (*Flk1+Etv2-Cdh5-*), differentiating (*Flk1+Etv2+Cdh5-*) and differentiated (*Flk1+Etv2-Cdh5+*) endothelial cell states in the E10.5 lung bud (**Figs. 4H-I**). Between the distally localized progenitors and proximally localized differentiated endothelial cells, we observed an enrichment of differentiating endothelial cells at the intersection of the two, suggesting their enrichment and subsequent incorporation at the forefront of existing networks (**Figs. 4J, Extended Data Fig. 7F**).

Altogether, these observations revealed a novel mode of vascularization that is distinct from both vasculogenesis and angiogenesis, which we call “integrative vasculogenesis”. In the posterior buds where *de novo* vascularization occurs, a subset of FLK1+ mesenchymal progenitors asynchronously commit to the endothelial fate and produce angioblasts. These angioblasts rapidly and randomly sample their environment as they differentiate. Upon the right maturation state and contact with a differentiating cell, newly differentiated cells are incorporated into existing networks. This continuous emergence of angioblasts and their intimate coordination with the existing vasculature is in stark contrast to classic vasculogenesis, which occurs in the absence of existing networks.

### Existing networks support the survival and patterning of newly differentiated endothelial cells

The above observation raised the possibility that existing networks play an important role in the incorporation of newly differentiated cells to the network. To test this, we performed a rescue experiment, in which differentiated endothelial cells in E10.5 Tg(*Flk1:GFP*) lung explants were chemically ablated and allowed to recover adjacent to an E12.5 Tg(*Cdh5Cre; lsl-tdTomato*) lung tissue with intact vascular networks^40,41^. Newly differentiated cells were GFP+, while rescue endothelial networks expressed tdTomato. After 24 hrs of co-culture post-ablation, a few tdTomato+ rescue networks invaded ablated tissues, suggesting that after drug treatment, the tissue was still able to support vascular development (**Fig. 5A**). Importantly, newly differentiated GFP+ endothelial cells also appeared, some of which integrated into the rescue networks (**Fig. 5A**). In contrast, far fewer GFP+ endothelial cells were recovered when explants were rescued with tissue lacking a tdTomato+ endothelial network (**Figs. 5B, C, Extended Data Figs. 9A-E**). Our result suggests that the existing vascular network increases the number of newly differentiated endothelial cells.

**Figure 5.**
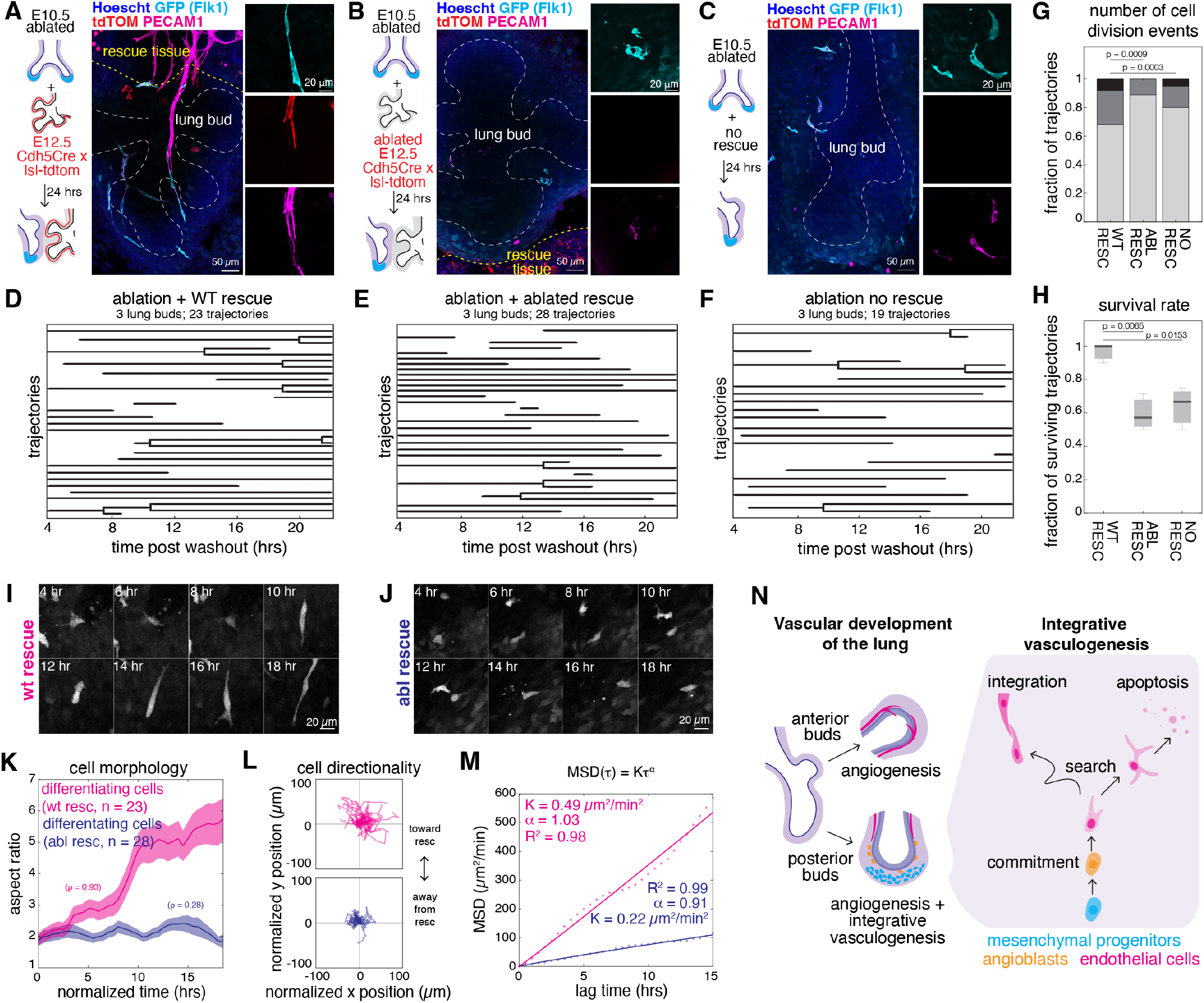
Existing networks support the survival and patterning of newly differentiated endothelial cells. **A-C**. Ablation-rescue experiments to test the role of existing endothelial networks in *de novo* vascularization. E10.5 Flk1-GFP explants underwent endothelial ablation for 24 hrs and then recovered for another 24 hrs in the presence of wild-type rescue tissue (**A**), rescue tissue with endothelial cells ablated (**B**), or no rescue tissue (**C**). Lungs from *Cdh5*^*+/Cre*^*;lsl-tdTomato* mice were used as rescue tissues, in which differentiated endothelial cells were tdTomato+. Explants were stained for GFP, tdTomato, and PECAM1. **D-F**. Lineage dynamics of all endothelial fate-committed cells tracked in the three conditions through timelapse imaging. **G-H**. Quantification of cell proliferation rates (**G**) and apoptotic events (**H**) based on the tracked lineages in **D-F**. n_wt-rescue_ = 23, n_abl-rescue_ = 28, and n_no-rescue_ = 19 traces across 3 experiments per condition. A two-sample t-test was performed between conditions to assess significance. **I-J**. Representative montage of newly differentiating endothelial cells in the presence of wild type (**I**) or ablated (**J**) rescue tissue. GFP channel only shown. **K**. Quantification of cell morphology changes during the recovery phase. Significance of the trend (ρ) was assessed by Spearman’s correlation. **L**. Overlay of all differentiating endothelial traces in the presence of wild type or ablated rescue tissues, normalized to the cell’s initial position at t_norm_ = 0 hrs. **M**. Mean squared displacement of committed endothelial trajectories. **N**. Model of embryonic lung vascularization through a combination of angiogenesis and integrative vasculogenesis. E10-E12.5 mouse lung harbors a *Wnt2+Flk1+* mesenchymal progenitor population that can produce endothelial cells *de novo* through intermediate *Etv2+* angioblasts in posterior buds. During differentiation, cells search locally for neighbors in a random walk manner before adhering to existing networks. In the absence of endothelial networks, differentiating cells have compromised motility and fail to find interacting partners, resulting in an abnormal morphology and in some cases apoptosis.

To ask whether the differential abundance of *de novo* generated GFP+ endothelial cells was due to the differences in progenitor fate commitment, or in the proliferation or survival of endothelial cells, we performed live imaging during recovery (**Supplementary Videos 3-5**). Immediately after chemical ablation, we observed scattered cells with higher levels of GFP (**Extended Data Figs. 9F, G**). Examination of cell states revealed these cells expressed little to no *Cdh5*, suggesting that despite the higher GFP level observed, these are newly differentiating endothelial cells that have yet to express the endothelial transcriptional profile (**Extended Data Fig. 9F, G**).

After washout, we observed a similar number of differentiating cells present across all conditions, likely due to the commitment of new endothelial cells during the ablation time window, which coincides with the peak period of angioblast emergence between E10.5-11.5 *in vivo* and during the first 24 hrs of *ex vivo* culture (**Figs. 5D-F, 2K, Extended Data Fig. 7B**). During the recovery phase, we observed a few rare FLK1_switch cells appearing shortly after washout, consistent with the drop off in endothelial commitment after E11.5 (**Figs. 5D-F, 2K**). Tracking these cells over the next 20 hrs, we observed that in the absence of existing networks, differentiating cells exhibited a reduction in the number of proliferation events (**Fig. 5G**). Moreover, while almost all tracked differentiating cells cultured with wild type explants survived until the end of the timelapse, approximately ⅓ of tracked cells underwent apoptosis in the absence of existing networks (**Fig. 5H**). This suggests that the presence of existing networks promotes the proliferation and survival of newly generated endothelial cells.

Furthermore, the search behavior of differentiating cells differed drastically across conditions. In the presence of wild type rescue networks, differentiating cells rapidly entered a migratory state to undergo a random search, similar to that observed in wild type explants (**Figs. 5L, M, Extended Data Fig. 10B, Supplementary Video 6**). Accompanying the increased motility, cells also adopted an elongated morphology (**Figs. 5I-K, Supplementary Video 6**). In contrast, differentiating cells in the absence of existing networks failed migrate, resulting in a much more restricted search space (**Figs. 5L, M, Extended Data Fig. 10B, Supplementary Video 7**). These cells also failed to adopt an elongated morphology (**Figs. 5I-K, Supplementary Video 7**). Importantly, these behaviors were unique to differentiating endothelial cells, as *Flk1+* mesenchymal progenitors maintained a rounded morphology, and randomly migrated at a slower speed (**Extended Data Figs. 10C-E**). These data suggest that mature networks may promote the search behavior of newly differentiated cells from the distal progenitor pool, even though the promoted migration effect does not appear directional.

Altogether, these data suggest the important role of existing networks in promoting the survival and motility of newly differentiating cells. In the absence of existing networks, newly differentiated cells exhibit abnormal morphologies, and a highly restricted search space before ultimately undergoing apoptosis. We speculate that in the absence of existing networks, newly differentiated cells are unable to form stable cell-cell contacts with like cells, impacting their survival.

## Discussion

In this study, we resolve a long-standing debate on the origin of pulmonary vasculature, by showing that the lung is vascularized through both local *de novo* endothelial specification and invasion from pre-existing vessels. We identify a *Wnt2+Flk1+* mesenchymal progenitor population in posterior lung buds that continuously generate *Etv2*+ angioblasts, which contribute directly to the pulmonary endothelium. In contrast, more anterior regions lack this progenitor pool and are vascularized through angiogenesis (**Extended Data Fig. 7G**). Thus, vascular assembly in the lung reflects the coordinated contribution by spatially distinct endothelial sources.

Mechanistically, we defined “integrative vasculogenesis” as a mode of vascular expansion that enables the incorporation of nascent endothelial cells into pre-existing networks (**Fig. 5N**). Differentiating endothelial cells adopt a highly migratory state and undergo exploratory movement before forming stable contacts with proximal vessels. Functional perturbation demonstrates that pre-existing networks are required to support the migration, proliferation and survival of nascent endothelial cells. In their absence, newly differentiated endothelial cells fail to adopt appropriate morphology and have elevated apoptosis, suggesting that network connectivity provides critical cues for endothelial integration and maturation. Similar phenomena have been observed in other adhesion-dependent epithelial tissues^42,43^. The molecular mechanisms mediating these interactions remain to be defined.

Finally, the identification of a mesenchymal progenitor source of endothelial cells has important implications for the generation of organotypic vasculature *in vitro*. Although a few specialized endothelial subtypes have been derived from pluripotent stem cells, recreating most organ-specific vascular networks remains a major challenge, in part due to limited understanding of their developmental origins^44,45^. Our finding suggests that endothelial cells with organ-specific identity may arise from lineage-restricted mesenchymal progenitors, raising the possibility that engineering such intermediate states could facilitate the generation of organotypic vasculature *in vitro*. Together, our findings establish integrative vasculogenesis as a mechanism that unifies distinct endothelial sources into a continuous vascular network, and has implications for the generation of functional vasculature in organoid models.

## Acknowledgments

We would like to thank Mark Greenwood, Diep Nguyen, Christina Lilliehook for their feedback on the manuscript, Chris Chen, Darrell Kotton and the rest of the Li Lab for helpful discussions. We would also like to thank Cassandra Rogers and Asier Marcos Vidal at the W.M. Keck imaging facility, and Jennifer Love, Sumeet Gupta and Stephan Mraz of the Whitehead Institute Genome technology core for their assistance. This work was supported by National Institute of Health grants DP2HD108777 (P.L.), Allen Family Philanthropies (P.L.), Eugene Bell Career Development Professorship (P.L.), MathWorks Graduate Fellowship (M.M. & M.L.), Ayla Gunner Prushansky fund (D.F), and R56 HL167937 (D.F).

## Author contributions

M.M. and P.L. conceived the study. M.M. and M.L. performed the scRNA-seq collection. P.C. and D.F. performed the *Wnt2* lineage tracing and embryo isolation. M.M. led the experimental work and computational analysis. M.M. and P.L. wrote the manuscript. P.L. supervised the work.

## Supplementary Materials for

## Methods

### Mouse husbandry

All mouse work was performed in accordance with Massachusetts Institute of Technology’s Institutional Animal Care and Use Committee (IACUC), and Children’s Hospital of Philadelphia Institutional Animal Care and Use Committee. Female mice of ages 6-12 weeks were used for timed matings with male mice 6 weeks of age (up to 6 months of age). Plugs were checked prior to noon the following day and considered to be E0.5 at that time. The following mouse strains were used: staining of wild-type lungs were performed in C57BL6/J (JAX #000664) and CD1 (Charles River, #022) mice. Live imaging and ablation-rescue experiments were performed by crossing Flk1-GFP males (Kdr^tm2.1Jrt^/J, JAX #017006)^38^ with CD1 females. Rescue tissues were generated by crossing Cdh5-Cre (JAX #006137)^41^ males with lsl-Tdtomato (JAX #007914)^40^ females. For lineage tracing experiments, the Wnt2^CreERT2^ allele^3^ and the Rosa26 fluorescence reporter, R26R^EYFP^(JAX #006148)^30^ were crossed to obtain embryos.

### scRNA-seq isolation

Embryos from C57BL6/J mice were isolated and staged based on morphological features. Samples were separated based on stage and dissociated with Dispase I (Stem Cell Technologies NC9995391) and Collagenase Type I (Sigma Aldrich SCR103) for 30 mins at 37 °C, agitating with a wide-bore pipette every 10 minutes. Cells were washed with DPBS + 10% FBS, before proceeding with the 10X CellPlex protocol. In brief: cells were incubated for 30 mins on ice with conjugated antibodies against CD45 (Thermo Fisher, 17-0451-82) and TER-119 (Thermo Fisher, 17-5921-82). Samples were spun down at 300 rcf for 5 mins at 4°C. Cells were barcoded by resuspending in 100 mL Multiplexing Oligo (10X Genomics, PN-1000261) for 5 mins at room temperature, before washing with DPBS + 10% FBS. Cells were counted and pooled to achieve an equivalent proportion of cells for each barcode. DAPI was added immediately before cell sorting as a live-dead stain. Blood and dead cells were depleted from the cell suspension by FACS, and the remaining cells were collected for library prep and sequencing as described in the 10X Genomics 3’ end assay protocol, v3.1.

### scRNA-seq quality control and processing

Raw reads were mapped to the mouse genome using CellRanger. Mapped reads were processed using Seurat^46^ (v5.0.0) in R (v4.2.1). Cells with fewer than 200 unique features were excluded. Cell cycle genes were regressed out to prevent clustering by cell cycle. Integration of different batches was performed by Reciprocal Principal Component Analysis (RPCA), and data was clustered using the top 20 principal components. Trajectories were inferred by diffusion map analysis^39^ using destiny (v3.12.0).

### Immunofluorescence

Samples were fixed in 4% PFA in PBS overnight at 4°C. Samples were then washed 3x for 10 mins in PBS at room temperature. For sections, samples were incubated in 30% sucrose in PBS, then embedded in OCT, flash-frozen in liquid nitrogen and sectioned. Sections were then fixed in 4% PFA in PBS for an additional 10 mins, before washing with PBS and permeabilizing in ice-cold methanol for 10 mins. Samples were then blocked in 1% BSA in PBS + 0.1% Tween-20 (PBSTw) for 30 mins, before incubating with primary antibodies for 3-4 hrs at room temperature or overnight at 4°C. Samples were washed 3x with PBSTw before incubating with secondary for 2-3 hrs at room temperature or overnight at 4°C. Samples were then washed 3x, one wash with Hoechst, before mounting for imaging in 70% glycerol in PBSTw. For whole mount staining, samples were dehydrated in MeOH for permeabilization and storage at -20°C. Samples were gradually rehydrated in PBS + 0.1% Tween-20 (PBSTw) on ice. Samples were blocked in 1% BSA in PBSTw for 2 hrs at room temperature, before incubating in primary antibody overnight at 4°C. Samples were washed 3x 2-3hrs with PBSTw at room temperature before incubating overnight with secondary antibodies at 4C. Samples were washed 3×1hr at room temperature, one wash with Hoechst, before mounting for imaging in 70% glycerol in PBSTw. Primary antibodies used are as follows: 1:200 CD31 (MEC13.3), 1:200 FLK1 (B.309.4), 1:500 GFP (A10262), 1:200 TDTOMATO, 1:200 ACTA2-488(1A4), 1:200 EPCAM (PA519832), 1:100 SOX17 (AF1924), 1:200 vWF(F3520) 1:200 EMCN (V.7C7), 1:200 CAR4 (AF2414), 1:200 NRP2 (AF567). All donkey secondary antibodies were used at 1:2000 dilution as and are as follows: aChicken-488 (A78948), aRat-594 (A-21209), aRat-647 (A78947), aRabbit-488 (A-21206), aRabbit-594 (A-21207), aRabbit-647 (A-31573), aRabbit-750 (SA000087), aGoat-488 (A-11055), aGoat-546 (A-11056), aGoat-647 (A-21447).

### HCR

All samples were processed with HCRv3 reagents using the Molecular Instruments protocol with slight modifications^47^. In short, samples were fixed 1% PFA in PBS overnight at 4°C or 4% PFA in PBS at room temperature for 30 mins. Samples were washed 3x for 10 mins with PBS + 1:1000 SUPERase inhibitor (AM2696), then dehydrated in MeOH for permeabilization and storage at -20°C. Samples were gradually rehydrated in PBSTw on ice. Samples were incubated for 30 mins at 37°C in pre-heated probe hybridization buffer, then incubated overnight at 37°C with probes (5nM probe final concentration). Samples were washed with 37°C probe wash buffer, then incubated overnight at room temperature in amplification buffer with 1:200 dilution of each hairpin. Samples were washed in 5xSSCT and mounted in 70% glycerol in 5xSSCT for imaging.

### Microscopy

All images were taken on a Zeiss LSM710 laser scanning confocal, or Leica Stellaris 8 white light laser scanning confocal. The following objectives were used on the 710: 10x air objective (NA 0.30), 40x oil objective (NA 1.3). The following objectives were used on the Stellaris: 40x glycerol (NA 1.25).

### Lineage tracing

Tamoxifen (Sigma-Aldrich, T5648) was prepared at 20 mg/mL by dissolving in a mixture of ethanol and corn oil, with a final v/v ratio of 10% ethanol and 90% corn oil. Recombination was induced in pregnant females by injecting tamoxifen via oral gavage at a concentration of 200 mg/kg of mouse. Immediately post tamoxifen oral gavage, we further performed intramuscular injection of medroxyprogesterone 17-acetate (Sigma-Aldrich, M1629) at a concentration of 2 mg/30 g of mouse to avoid preterm birth.

### Cell segmentation and classification of fixed images

Cellpose-SAM^48^ was used to segment nuclei by Hoechst staining. Masks from individual optical sections were pooled and analyzed to determine the proportion of cell states in MATLAB. Signal thresholds were determined empirically by negative regions in the tissue, and were used to classify cells by expression of a given transcript or protein.

### Explant culture

Embryonic lungs were isolated and cultured in Advanced DMEM/F12 (Life Technologies, 12634010) supplemented with 5% stem-cell grade FBS (Life Technologies, 10439024) and Penn/Strep (Life Technologies, 15140122). Explants for live imaging were plated in CellVis 24-well glass bottom imaging plates (CellVis, P24-1.5H-N) coated with rat tail collagen (Life Technologies, A1048301). Explants were cultured for 12 hrs in the incubator with 150 mL of media to prevent explant floating from the coverslip. Once adhered, wells were supplemented with additional warmed media (300 mL total) before starting imaging. All empty wells were filled with water to avoid evaporation. Explants were imaged on the Leica Stellaris 8 white light laser scanning confocal, with a 40x glycerol objective (NA 1.25) at 0.75x magnification. Z-stacks of 2μm thickness were taken at 20-30 min intervals for 24-36 hrs.

### Explant ablation and rescue experiments

Endothelial cells were ablated using 10 μM SU5416 (R & D Systems, 3037) in explant media. Once ablated, explants were washed 3X with media to remove any residual drug. Explants were transferred to a well of a 24-well glass bottom imaging plate and incubated for an additional 24 hrs in explant media for live imaging and/or analysis. For rescue tissues, E12.5 lungs were isolated from a Cdh5-Cre x lsl-tdTomato timed mating. Rescue lungs were cut into small pieces and plated onto a collagen-coated 24-well glass bottom imaging plate for 24hrs before plating ablated E10.5 tissues adjacent to rescue tissues. A 24hr dose of 5 μM Ki8751 (Cayman Chemical Company, 18004-10) was used to ablate rescue tissues. Ablated rescue tissues were washed 3X with media before plating ablated E10.5 tissue.

### Cell segmentation and analysis of live images

Analysis of timelapse imaging was performed on maximum projected z-stacks. For FLK1_low and FLK1_high cells, a Cellpose model was trained to recognize the diverse cell morphologies from GFP+ cells. This model was used in trackmate^49^ (v7.11.1) to detect cells. The Linear Assignment Problem (LAP) algorithm^50^ was used to link cells into trajectories, with distance and gap closing thresholds set to 30 μm, penalizing linkages with drastically different cell size and mean intensities between consecutive frames. FLK1_switch masks were generated using a separate Cellpose model. Masks were manually evaluated and corrected if necessary. The mask output from the segmentation was then fed into TrackMate for LabelMap detection and LAP trajectory assignment. Trajectories were manually corrected in instances of erroneous linkages. For all populations, trajectories with fewer than 10 spots were discarded. All trajectories and masks were then imported into MATLAB for further analysis. By virtue of the segmentation approach above, FLK1_switch cells were already classified. FLK1_high and FLK1_low cells were classified simply by GFP expression levels, determined by mean GFP intensity over the course of the timelapse, with FLK1_high cells assigned if the intensity exceeded that of the average FLK1_low cell.

For ablation-wash out experiments, differentiating cells were segmented as the FLK1_switch population: Cellpose segmentation and manual correction, followed by TrackMate LabelMap detection and LAP trajectory assignment. Masks and trajectories were imported into MATALB for further analysis.

### Quantification of motility dynamics

The instantaneous speed to each trajectory at a given time was calculated as follows:

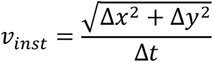

Where Δx and Δy are the difference in x, y positions between two consecutive spots in a trajectory, with Δt as the difference in time between two consecutive spots in a trajectory.

The mean squared displacement for each trajectory was calculated as follows.

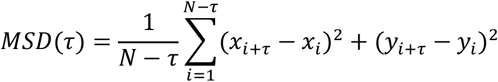

Where t is the time lag between frames, N is the total number of positions and x_*i*_,y_*i*_ are the coordinates at a given time step *i*.

To determine the diffusivity behavior, MSDs were fit to a power law as follows using the curve-fitting toolbox in MATLAB:

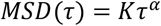

Where a designates the type of diffusion, with α <1 being sub-diffusive, α = 1 as random motion, and α > 1 as supra-diffusive. *K* is interpreted as an effective diffusion coefficient.

**Extended Data Figure 1.**
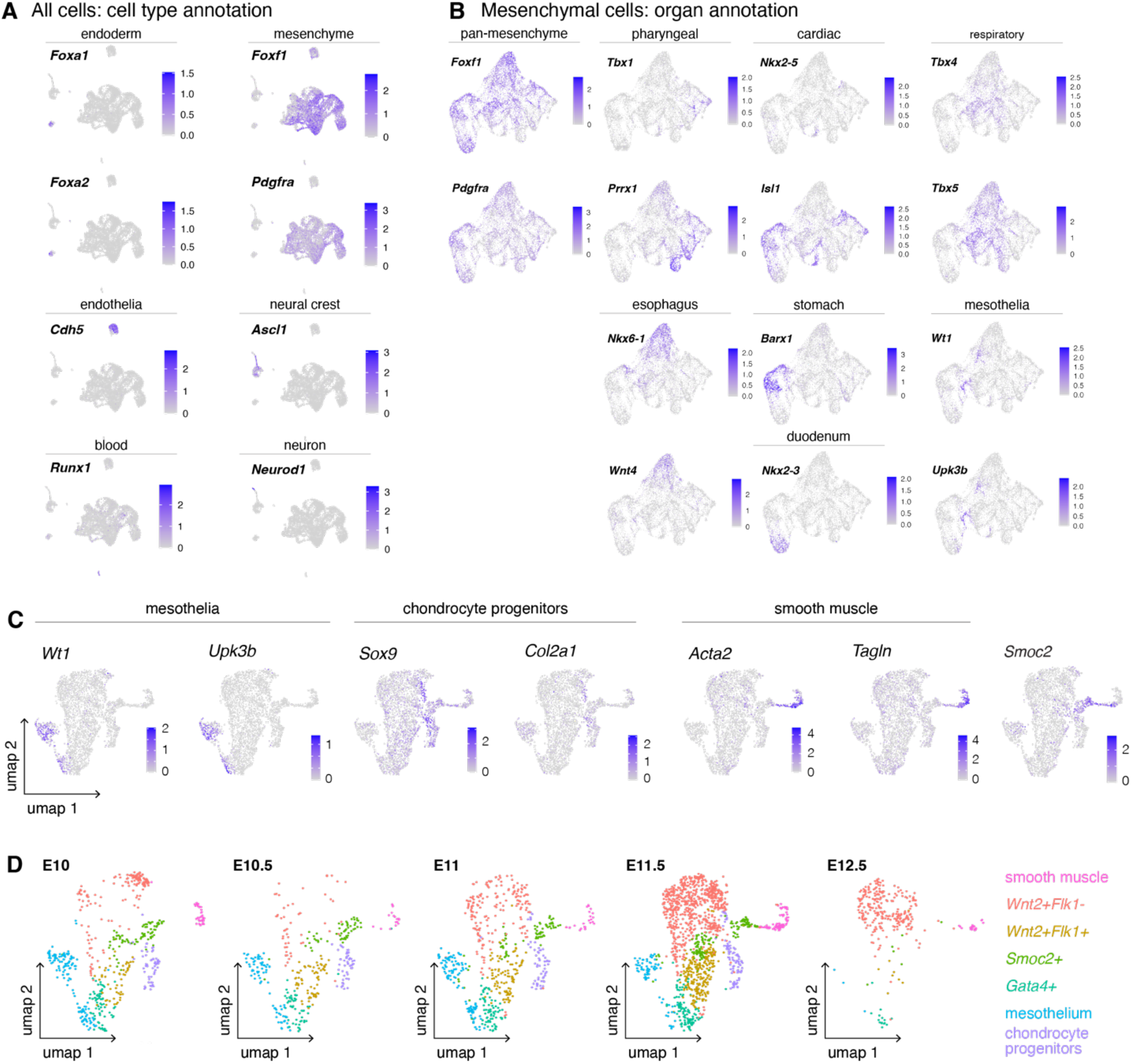
Annotation of scRNA-seq data. **A**. Cell type annotation of the entire dataset as follows: endoderm (*Foxa1+Foxa2+*), mesenchyme (*Foxf1* and/or *Pdgfra*), blood (*Runx1*), endothelia (*Cdh5*), neural crest (*Ascl1*) and neurons (*Neurod1*). **B**. Organ identities were defined as follows: pharyngeal (*Tbx1+*/*Prrx1+*/*Isl1+*), cardiac (*Isl1+*/*Nkx2*.*5+*), respiratory (*Tbx4+Tbx5+*), esophagus (*Nkx6*.*1+Wnt4+*), stomach (*Barx1+*), duodenum (*Nkx2*.*3+*) and mesothelium (*Wt1+Upk3b+*). **C**. Expression of marker genes reflecting committed cell states in the respiratory mesenchyme. *Smoc2* may reflect a progenitor or subpopulation of smooth muscle cells. **D**. UMAP representation of respiratory mesenchymal cell populations split by stage.

**Extended Data Figure 2.**
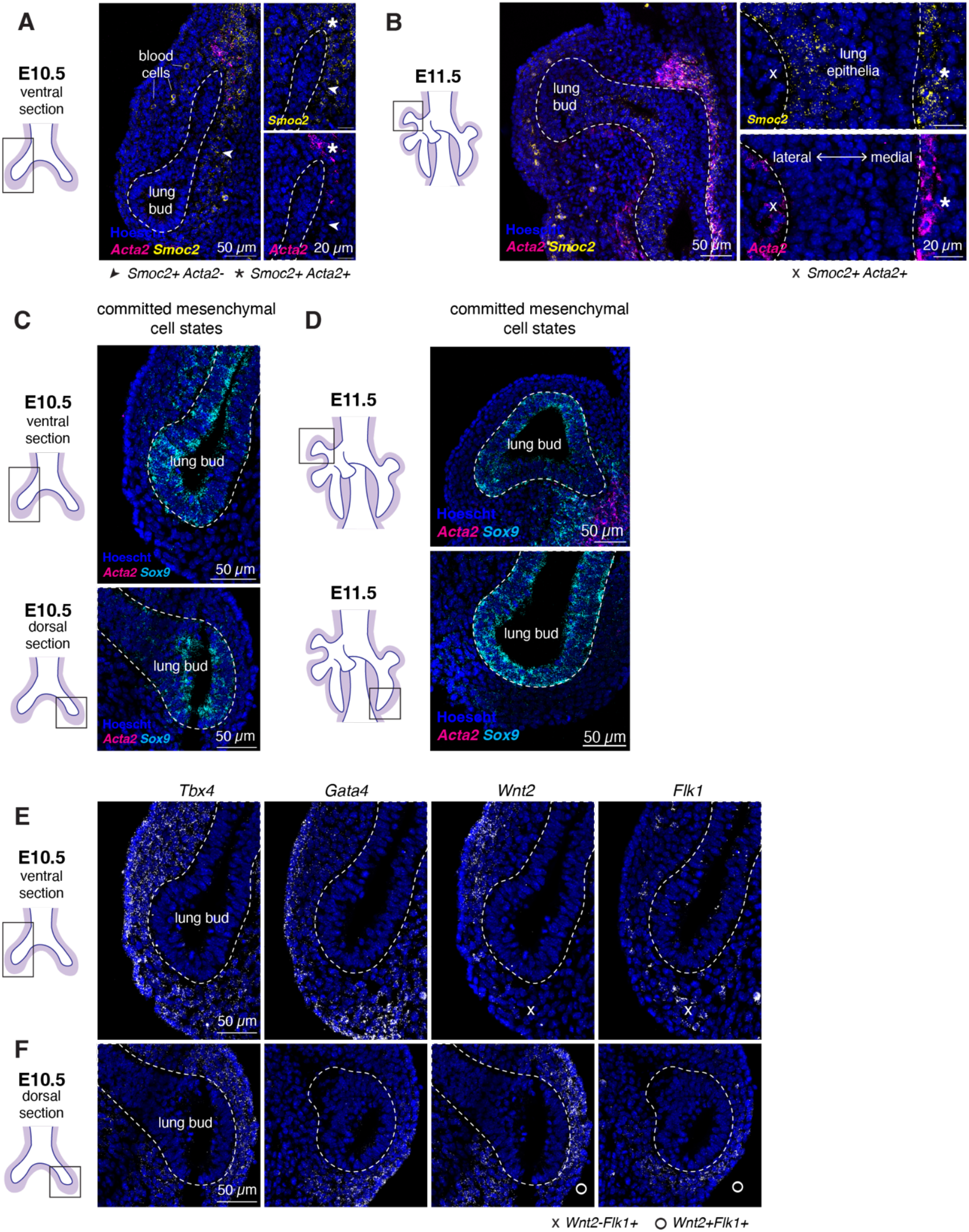
Spatiotemporal distribution of mesenchymal cell states. **A-B**. HCR validation of *Smoc2* expression in the respiratory mesenchyme in E10.5 (**A**) and E11.5 (**B**) lung tissue sections. Curiously, medial smooth muscle cells maintained Smoc2 expression, while lateral smooth muscle cells did not (**B**). **C-D**. HCR visualization of committed respiratory mesenchymal populations in E10.5 (**C**) and E11.5 (**D**) tissue sections. *Sox9+* in the mesenchyme marks chondrocyte progenitors, *Acta2* marks smooth muscle cells. **E-F**. Validation of identified respiratory mesenchymal progenitors at E10.5. The top row represents a more ventral section of the E10.5 lung, while the bottom row represents a more dorsal section.

**Extended Data Figure 3.**
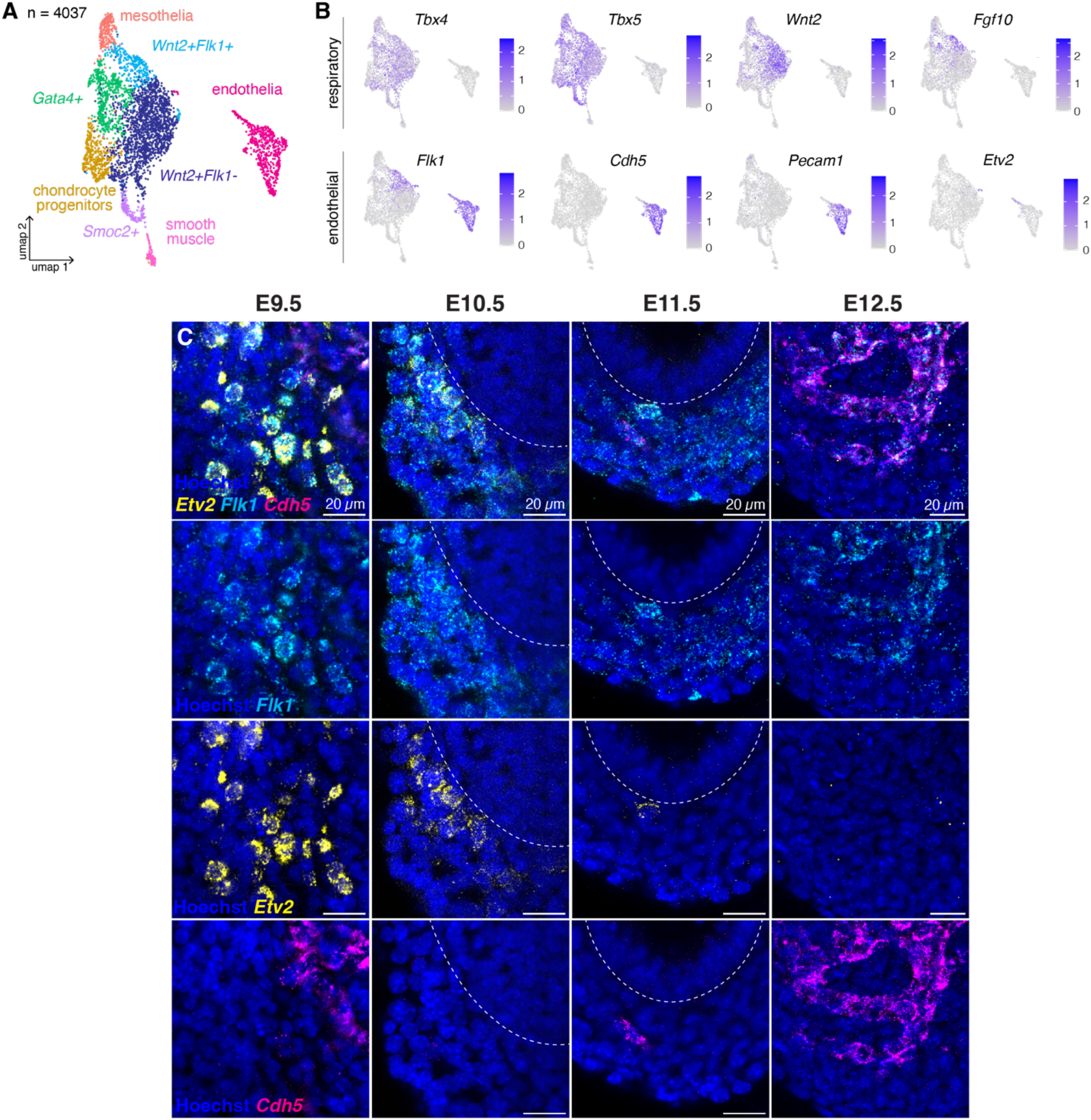
Characterization of heterogeneous FLK1+ populations in the lung. **A**. UMAP representation of respiratory mesenchymal cells and endothelial cells from single cell isolation. **B**. Expression of respiratory marker genes (top row) and endothelial marker genes (bottom row) in mesenchymal and endothelial clusters. **C**. HCR endothelial cell state markers. *Etv2+* cells are *Cdh5-* and remain in the *Flk1+* domain. *Etv2* channel is shown in Figure 2 E-H.

**Extended Data Figure 4.**
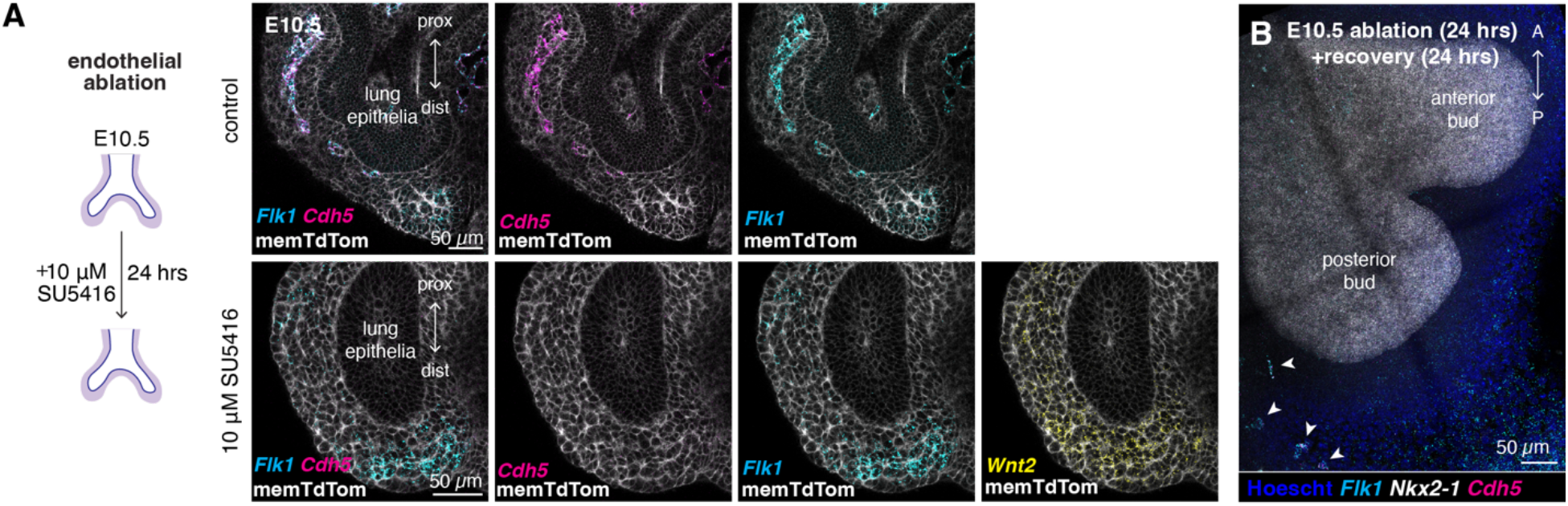
Validation of endothelial ablation by SU5416. **A**. Chemical ablation validation after 24 hrs incubation with VEGFR2 inhibitor SU5416. Endothelial cells (*Flk1+Cdh5+*) are absent but *Wnt2+Flk1+Cdh5-* progenitors remain post ablation. **B**. E10.5 lung bud post ablation and recovery. Anterior bud in the field is devoid of any new endothelial cells.

**Extended Data Figure 5.**
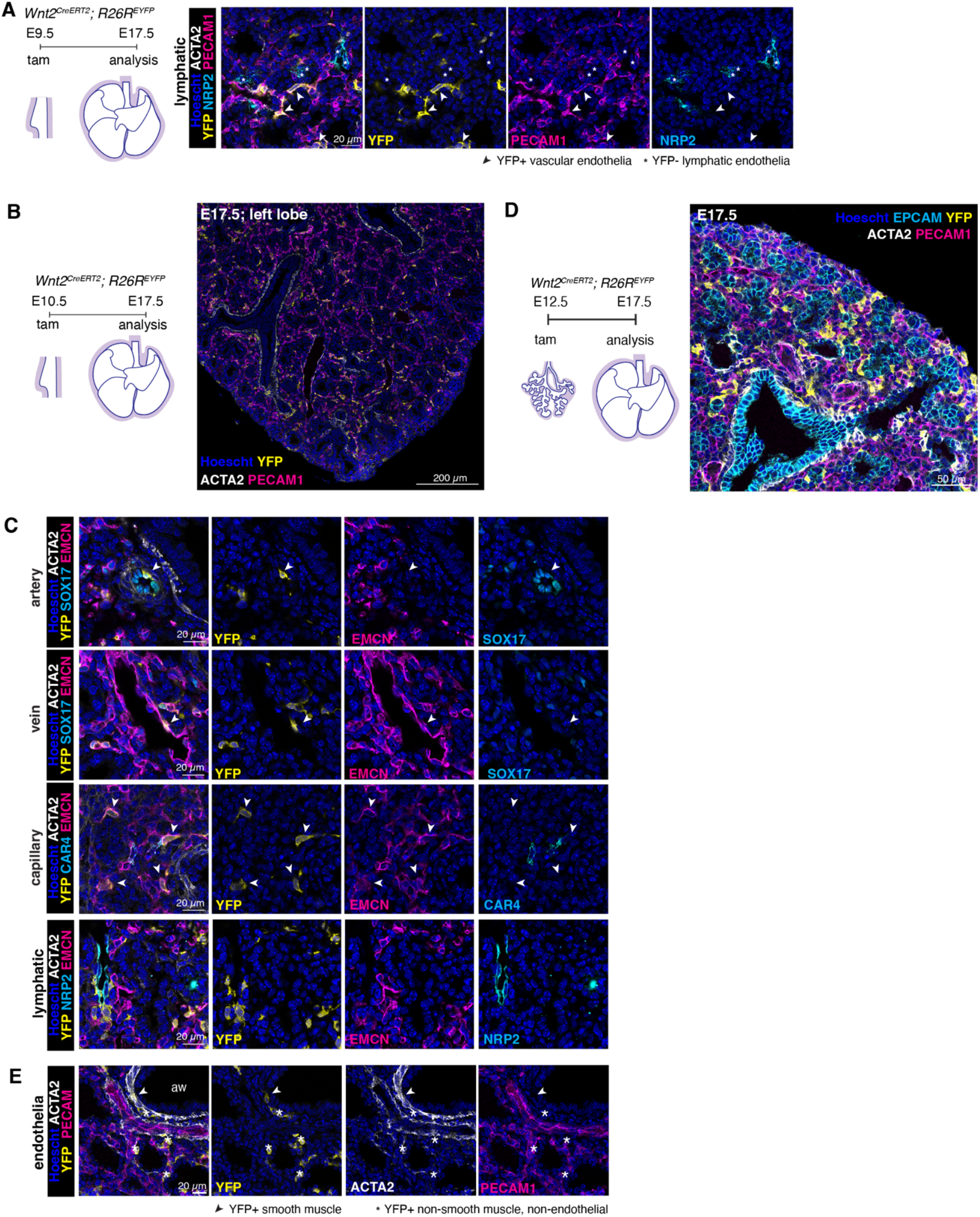
*Wnt2* lineage tracing labels pulmonary endothelium prior to E12.5. **A**. Representative tissue section of E17.5 mouse lung from lineage tracing experiments, in which tamoxifen was administered at E9.5. NRP2+ lymphatic endothelium was not observed in the samples examined. **B-C**. E10.5 *Wnt2* lineage tracing. Representative tissue section of E17.5 mouse lung from lineage tracing experiments, in which tamoxifen was administered at E10.5 (**B**). Artery, vein and capillary endothelial cells were labelled, whereas lymphatic endothelium was not labelled (**C). D-E**. E12.5 *Wnt2* lineage tracing. Representative tissue section of E17.5 mouse lung from lineage tracing experiments, in which tamoxifen was administered at E12.5 (**D**). Of the samples examined, no endothelial cells were labelled, despite labelling of adjacent stromal cells (**E**).

**Extended Data Figure 6.**
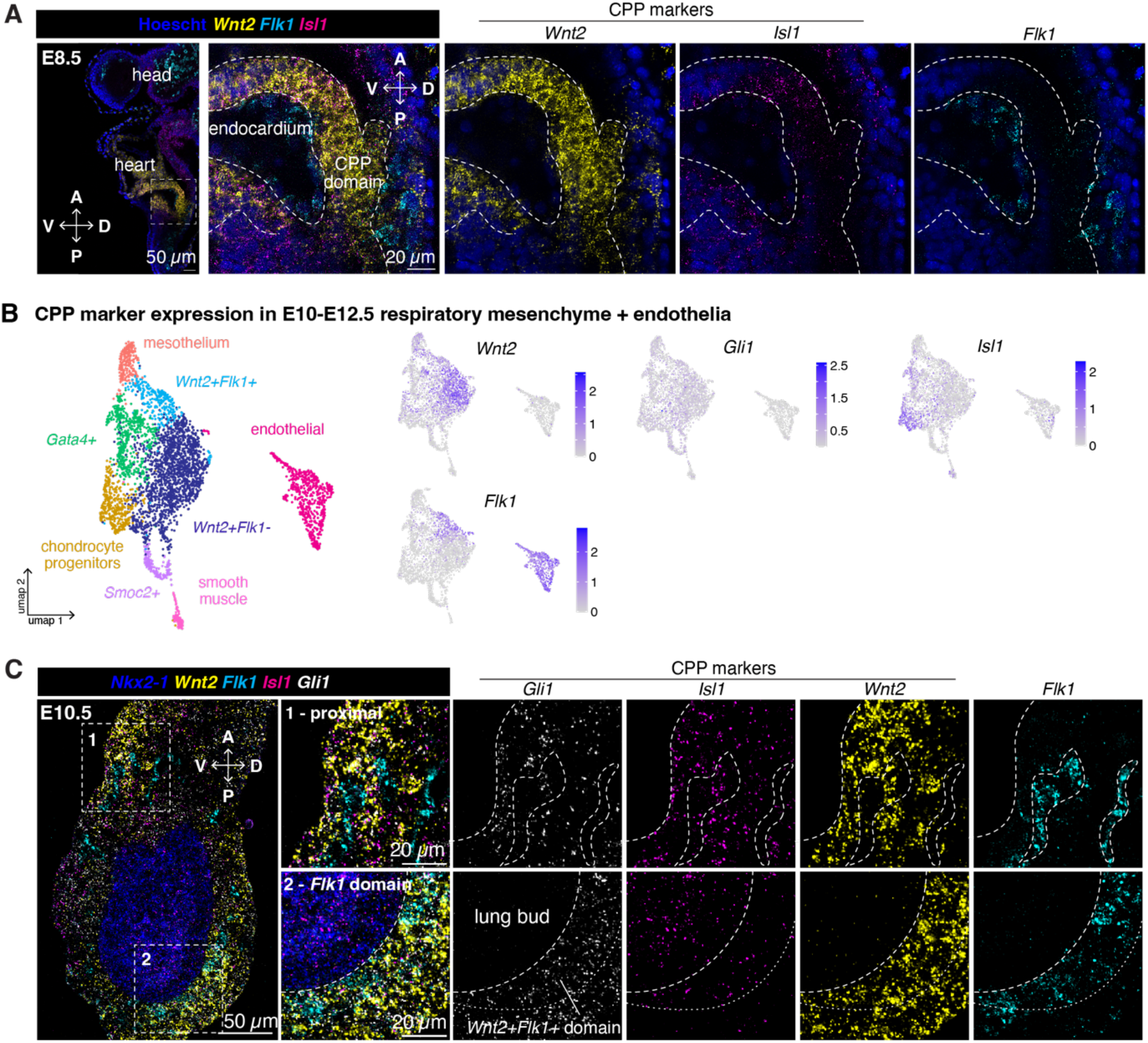
*Wnt2+Flk1+* progenitors are spatially and temporally distinct from CPPs. **A**. HCR of E8.5 anterior foregut. The CPP domain, defined by the co-expression of *Wnt2* and *Isl1*, is devoid of *Flk1* expression. **B**. Expression of CPP markers in our E10-E12.5 scRNA-seq dataset. *Isl1* and *Flk1* expression were largely mutually exclusive. **C**. Optical section of an E10.5 lung, probing for *Flk1* and CPP markers *Wnt2, Gli1* and *Isl1*. Ventrally, a *Wnt2+* population expressing higher levels of *Isl1*, resembling CPPs, does not express *Flk1*. Conversely, dorsally and in the posterior region of the organ, *Wnt2+* cells which express *Flk1* have little to no *Isl1* expression in this domain. As Sonic hedgehog (SHH) signaling is essential for lung development^51,52^, all mesenchymal cells express the SHH response gene *Gli1*.

**Extended Data Figure 7.**
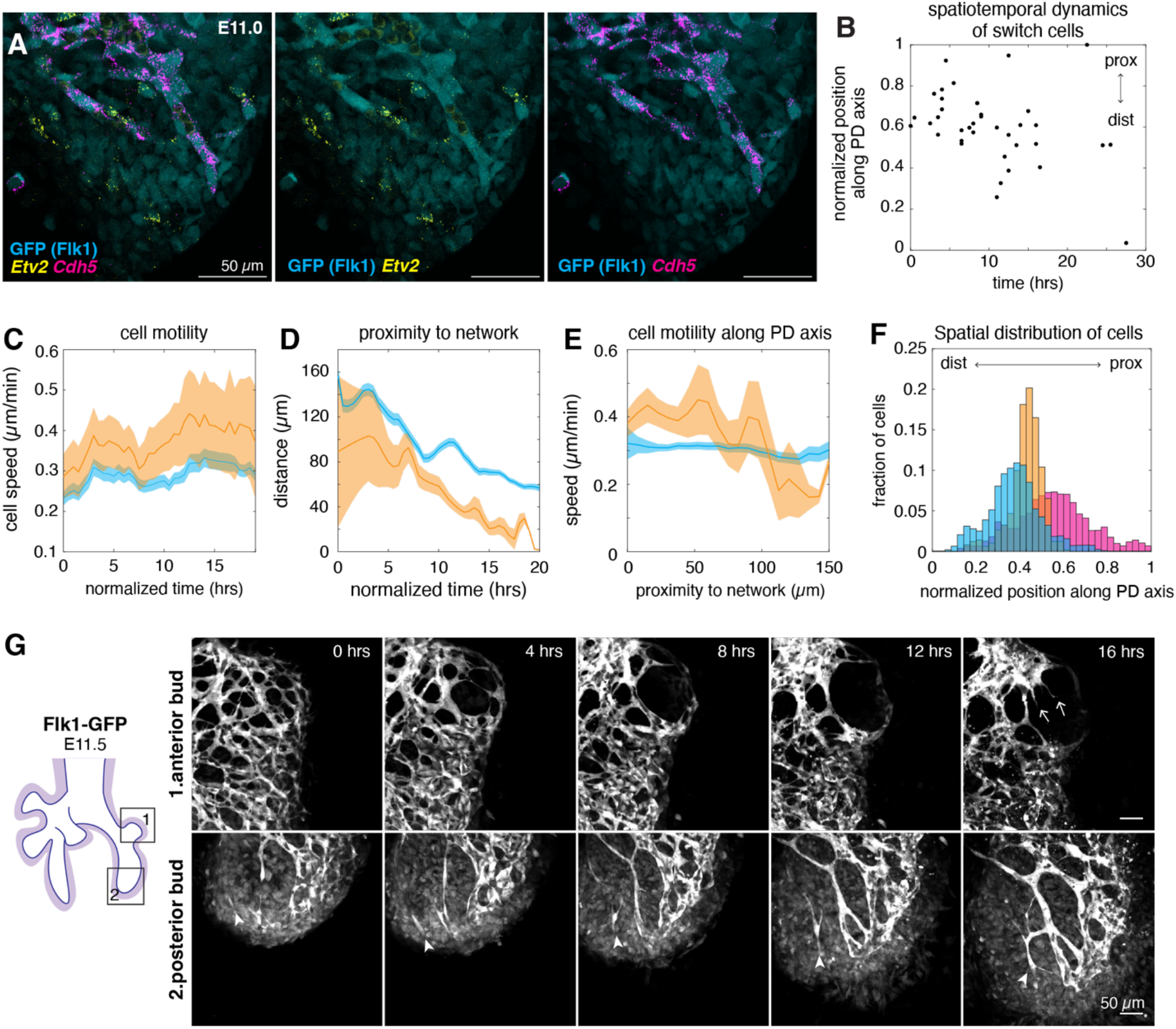
FLK1-GFP cell states and dynamics in explants. **A**. Representative images of *Etv2* and *Cdh5* expression in FLK1-GFP E11 lung buds. The levels of GFP protein and *Etv2/Cdh5* transcripts could be quantified for the same cells, enabling the establishment of correlation between GFP intensity and cell states. **B**. Temporal emergence of FLK1_switch cells during timelapse. 38 cells pooled over 4 independent samples. Position along the proximal-distal axis is normalized within the FLK1_switch population, with cells nearest the proximal networks at a normalized position of 1, and cells furthest at position 0. **C**. Cell speed over time, pooled across all samples. The mean speed and SEM was calculated for each frame during the timelapse. **D**. The mean and SEM proximity of FLK1_switch and FLK1_low cell states to the nearest FLK1_high cell over time. **E**. FLK1_switch and FLK1_low cell speeds as a function of distance to nearest FLK1_high cell. The mean and SEM is calculated for each frame. **F**. Spatial distribution of FLK1+ populations across all frames along the proximal-distal axis. **G**. Montage of an E11.5 explant timelapse. The top row represents the anterior bud of the left lobe, in which FLK1+ progenitors are absent. Networks appear to expand by angiogenesis, by the presence of tip cells extending into the avascular region (arrow). The bottom row represents the posterior bud of the left lobe, with FLK1+ progenitor domain present (arrowhead).

**Extended Data Figure 8.**
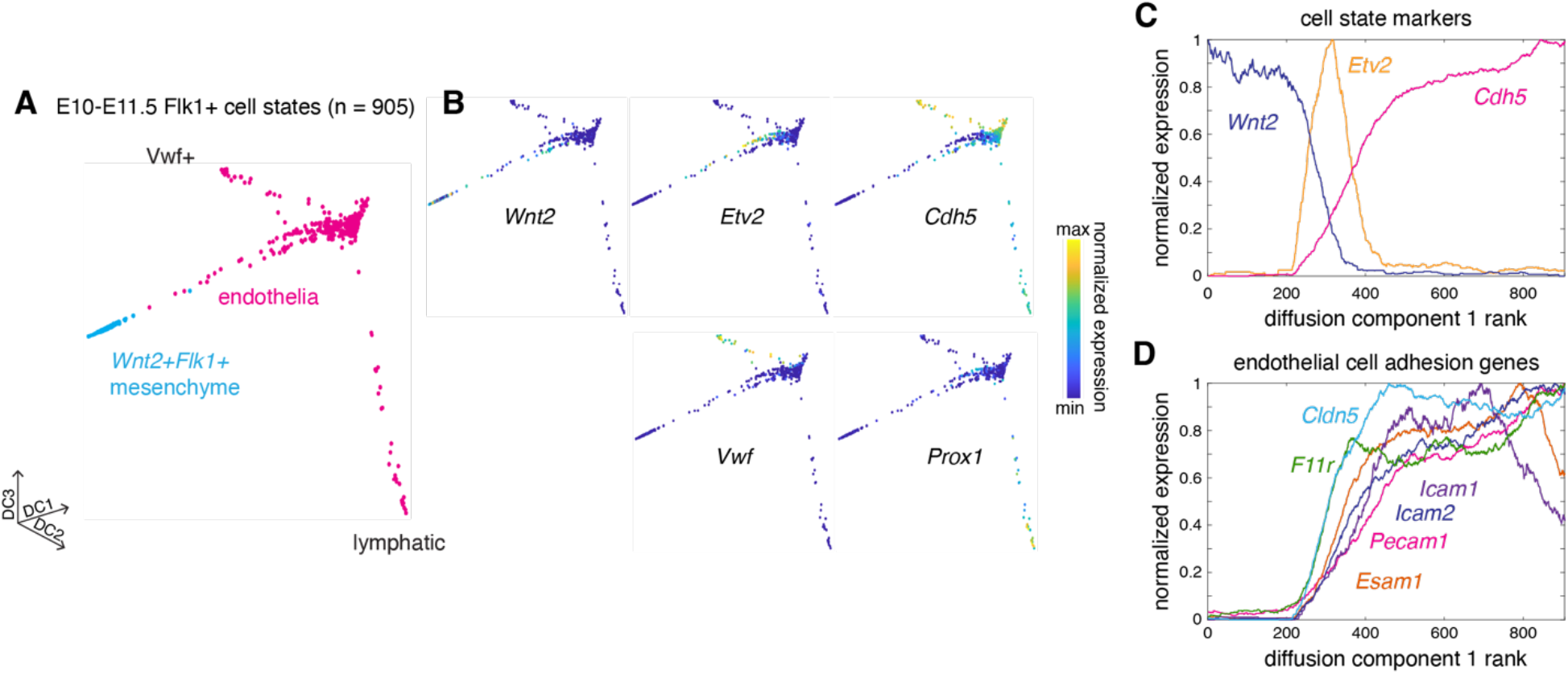
Transcriptional changes in differentiating endothelial cells. **A-B**. Diffusion map^39^ representation of *Wnt2+Flk1+* mesenchymal progenitors and endothelial cells. Marker gene expression along the different diffusion components. Cells which deviate from the endothelial lineage represent more differentiated *Vwf+* endothelial cells, or *Prox1+* lymphatic endothelium. **C**. Cell state markers along diffusion component 1, which recapitulates the expected differentiation trajectory from *Wnt2+Flk1+* progenitors to endothelial cells. **D**. Expression of cell adhesion molecules during endothelial differentiation.

**Extended Data Figure 9.**
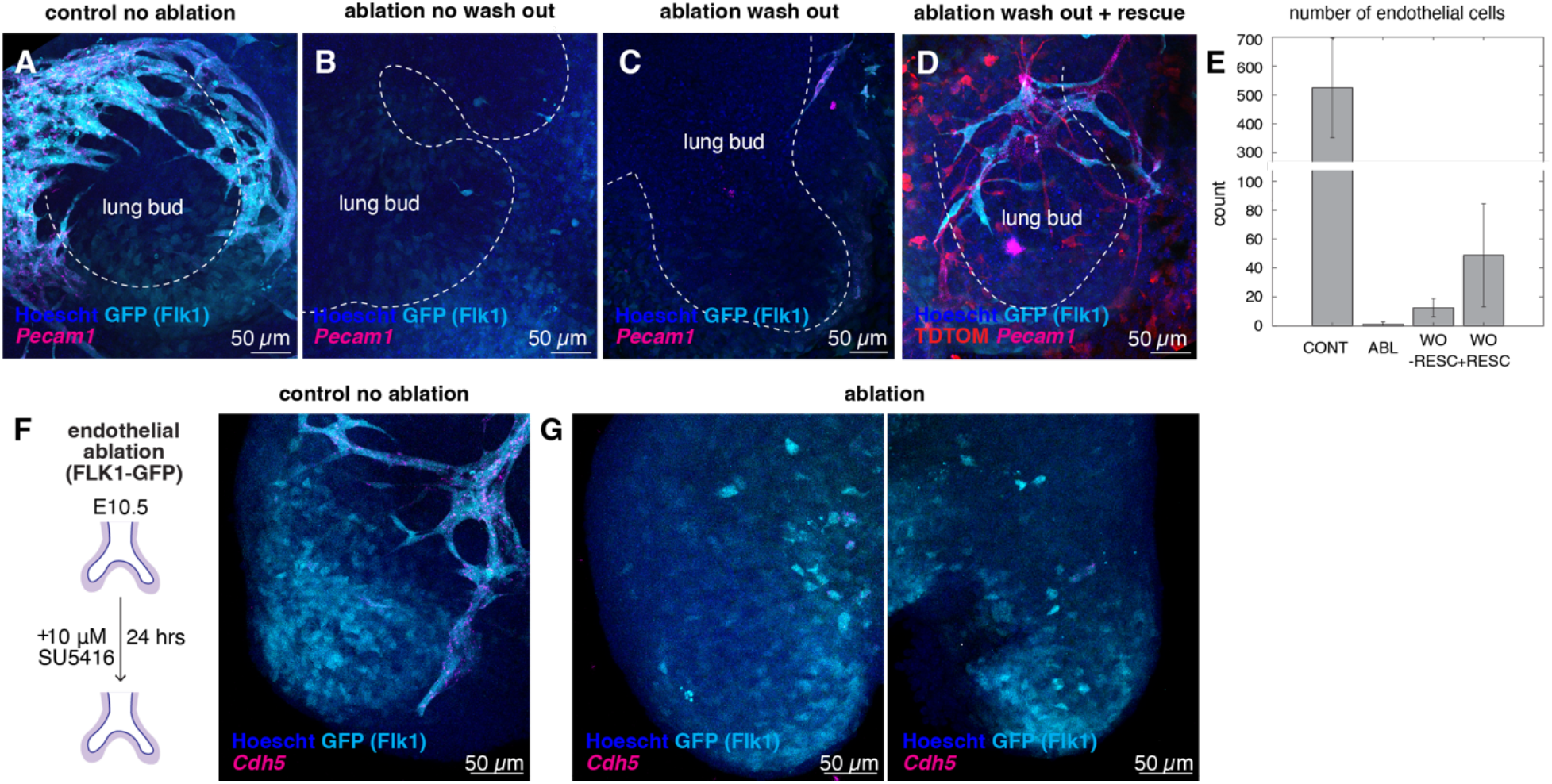
Endothelial number is increased in the presence of existing networks. **A-D**. Representative maximum intensity projection images of E10.5 Flk1-GFP explants after 48 hrs of culture (**A**), 48 hrs of culture in SU5416 (**B**), 24 hrs of recovery post chemical ablation (**C**) and 24 hrs of recovery post chemical ablation, in the presence of a wild type E12.5 rescue tissue (**D**). **E**. Quantification of the number of GFP+ endothelial cells post ablation and recovery experiments. 3 independent lung explants were quantified per condition. Mean and standard deviation are shown. **F-G**. Chemical ablation validation after 24 hrs incubation with VEGFR2 inhibitor SU5416 in FLK1-GFP embryos. GFP+ cells express *Cdh5* in control explants (**F**) but not in ablated (**G**) explants.

**Extended Data Figure 10.**
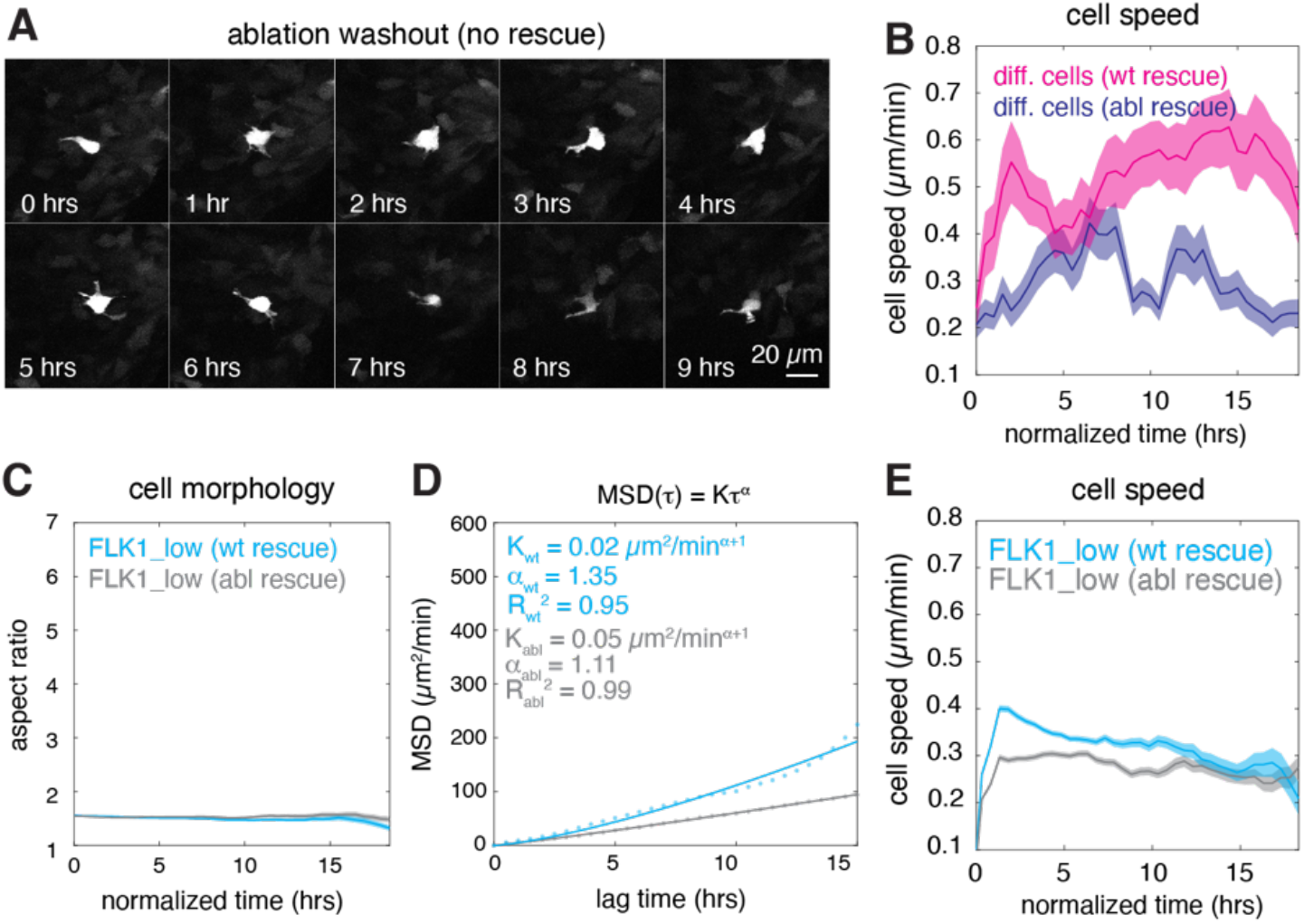
Dynamics of FLK1-GFP cells across ablation and rescue conditions. **A**. Representative montage of newly differentiating endothelial cells in the absence of any rescue tissue. **B**. Quantification of cell speed over time, pooled across 3 independent samples for each condition. Mean and SEM plotted. **C**. Cell morphology dynamics of FLK1_low (mesenchymal progenitors) in ablation recovery experiments. n_wt_ = 1040 trajectories. n_abl_ = 849 trajectories. Mean ± SEM. **D**. Mean squared displacement of FLK1+ mesenchymal progenitors. **E**. Quantification of cell speed of FLK1_low cells over time. Mean ± SEM.

## Supplementary Videos

**Supplementary Video 1. Incorporation of newly differentiated endothelial cells into existing networks**. Maximum intensity projection of FLK1-GFP explants imaged on Leica Stellaris laser scanning confocal. Stacks were acquired every 20 minutes at 2 μm z intervals. Montage from this movie is shown in Figure 4A.

**Supplementary Video 2. Cell morphology dynamics of FLK1_switch cells**. Example FLK1_switch cell as it upregulates its GFP expression an undergoes dynamic changes in cell shape before stabilizing. Montage from this movie is shown in Figure 4D.

**Supplementary Video 3. Differentiating endothelial cell dynamics in the presence of wild-type rescue networks**. Time-lapse video of ablation-recovery in the presence of wild-type rescue tissue with tdTomato+ endothelial networks. Differentiating cells are outlined in white. tdTomato+ rescue networks are in magenta. Analysis is shown in Figure 5D-H.

**Supplementary Video 4. Differentiating endothelial cell dynamics in the presence of ablated rescue networks**. Time-lapse video of ablation-recovery in the presence of ablated rescue tissue. Differentiating cells are outlined in white. tdTomato+ blood cells derived from hemogenic endothelium are visible. Analysis is shown in Figure 5D-H.

**Supplementary Video 5. Differentiating endothelial cell dynamics during ablation recovery**. Time-lapse video of ablation-recovery in the absence of any rescue tissue. Analysis is shown in Figure 5D-H.

**Supplementary Video 6. Motility and migration of differentiating endothelial cells in the presence of WT rescue networks**. Example of a newly differentiating cell after ablation in the presence of WT rescue networks. Analysis is shown in 5K.

**Supplementary Video 7. Motility and migration of differentiating endothelial cells in the presence of ablated rescue networks**. Example of a newly differentiating cell after ablation in the presence of ablated rescue networks. Montage from this movie is shown in Figure 5J.

